# Pioglitazone administration restores a PPARG-dependent transcriptional network and ATP levels within skeletal muscles of mice implanted with patient-derived breast tumors

**DOI:** 10.1101/2021.10.04.463044

**Authors:** David A. Stanton, Hannah E. Wilson, Matthew G. Chapa, Jessica N. Link, Werner Geldenhuys, Emidio E. Pistilli

## Abstract

**Background:** Fatigue is common in patents with breast cancer (BC), and can occur in patients with early stage disease and in the absence of muscle wasting (i.e. cachexia). We have reported transcriptional and proteomic alterations in muscles from BC patients, which are associated with fatigue. Mice implanted with human BC xenografts recapitulate the muscle molecular composition changes seen in patients, coupled with a greater rate of contraction-induced fatigue. Multiple bioinformatics platforms in both human and mouse muscles have identified peroxisome proliferator activated receptor gamma (PPARG) as central to this phenotype, with multiple PPARG target genes downregulated in response to tumor growth. The current study tested the hypothesis that the PPARG agonist pioglitazone (pio), a commonly prescribed diabetes drug, would rescue the transcriptional alterations observed in muscles of tumor-bearing mice.

**Methods:** Sixteen female NSG mice were implanted with breast cancer patient-derived orthotopic xenografts (BC-PDOX) via transplantation of Her^2^/neu^+^ human tumor fragments. BC-PDOX mice were randomly assigned to a treatment group that received daily oral pio at 30 mg.kg^-1^ (n=8), or a control group that received a similar volume of vehicle (n=8). Treatment was initiated when tumors reached a volume of 600mm^3^, and lasted for 2-weeks. Hindlimb muscles were isolated from BC-PDOX and non-tumor bearing mice for RNA-sequencing, gene expression validation, and ATP quantification. Differentially expressed genes (DEGs) in muscles from BC-PDOX mice relative to non-tumor bearing controls were identified using DESeq2, and multiple bioinformatics platforms were employed to contextualize the DEGs.

**Results:** We found that the administration of pio restored the muscle gene expression patterns of BC-PDOX mice to a profile resembling muscles of non-tumor bearing NSG control mice. Validation of skeletal muscle gene expression by qPCR confirmed pio increased the expression of PPARG target genes in skeletal muscles. Isolated mitochondria from muscles of BC-PDOX mice treated with pio contained greater levels of ATP. There were no differences in body weights, muscle weights, or tumor volumes in pio vs. vehicle treated BC-PDOX mice.

**Conclusions:** These data demonstrate that oral pio supplementation rescues the BC-associated downregulation of PPARG target genes in skeletal muscle. Additionally, muscles from BC-PDOX mice treated with pio had greater levels of ATP, which would be associated a more fatigue-resistant muscle phenotype. Therefore, we propose that the FDA-approved and generic diabetes drug, pio, be considered as a supportive therapy for the treatment of BC-associated muscle fatigue.

## Introduction

Breast cancer (BC) is the most commonly diagnosed cancer and the second leading cause of cancer-related death in women^1^. Through increased medical surveillance, the rate of BC-associated deaths has decreased, especially in women over the age of 50. Greater medical surveillance and early detection of breast tumors may also be associated with the observation that BC-patients have a lower prevalence of diagnosed cachexia, defined as weight loss of >5% body weight, compared to multiple cancer types^2–4^. Despite the effectiveness of primary treatments, high 5-year survival rates, and low prevalence of cachexia, BC patients commonly report debilitating physical impairments including muscle fatigue. In particular, fatigue is reported by a majority of breast cancer patients, with up to 96% of all BC patients experiencing this symptom^5–7^. Incidence of fatigue contributes to adverse patient outcomes including dose reduction or treatment cessation of chemotherapy and radiation, increased pain and reduced quality of life, and higher rates of cancer recurrence^8–10^. Additionally, there are currently no effective drug therapies that target cancer-associated fatigue, or the larger cancer cachexia syndrome ^11,12^.

While BC may not be associated with skeletal muscle wasting in the majority of patients, recent data suggest that breast tumor growth does influence mitochondrial and metabolic pathways within skeletal muscle that are associated with fatigue^13,14^. Recent data from our laboratory in muscle biopsies obtained from BC patients as well as skeletal muscle collected from mice implanted with patient-derived breast tumors (BC-PDOX) demonstrate aberrant skeletal muscle gene expression patterns, skeletal muscle mitochondrial dysfunction, and BC-associated fatigue^13–15^. These BC-induced skeletal muscle intrinsic changes are all potential mechanisms for the fatigue observed in BC patients. Therefore, developing therapies specific to this skeletal muscle dysfunction may represent an effective strategy, which would be applicable to the many patients experiencing BC-associated fatigue. Analysis of transcriptional networks indicated that the BC-induced downregulation of the transcription factor peroxisome proliferator-activated receptor gamma (PPARG) was central to this phenotype within the skeletal muscle of both BC patients and BC-PDOX mice^13–15^. This was of interest as PPARG is a transcription factor that aids in whole body energy homeostasis via the regulation of genes involved in lipid metabolism and mitochondrial function^16–18^. Analysis of these skeletal muscle transcriptional changes strongly indicated that PPARG is the central regulator of pathways that contribute to the increased muscle fatigue experienced by patients and BC-PDOX mice^13–15^. In support of this analysis, we have recently observed that mitochondrial respiration is impaired and mitochondrial ATP content is decreased in mitochondria isolated from the skeletal muscle of BC patients and BC-PDOX mice^14,15^. Importantly, exogenous PPARG protein expression was able to ameliorate these effects in myogenic cultures *in vitro*^15^.

To address the potential BC-associated skeletal muscle mitochondrial dysfunction, we generated BC-PDOX mice with HER^2^/neu^+^ overexpressing tumors and supplemented them with the PPARG agonist pioglitazone (pio) for two weeks. We hypothesized that pio supplementation would increase the expression of PPARG target genes within skeletal muscle, thereby restoring PPARG transcriptional networks and improving mitochondrial function and ATP production. The data presented herein support a mechanism whereby pio restores PPARG transcriptional networks, resulting in improved skeletal muscle ATP content in BC-PDOX mice. This restoration of ATP content may reflect a rescue of mitochondrial function in muscle, thereby decreasing the fatigue experienced by BC patients. Therefore, pio supplementation in BC patients represent a supportive therapy that improves overall quality of life by reducing fatigue experienced by these patients. This reduction in fatigue, in turn, would allow patients to continue primary tumor directed therapies when they otherwise may not be able to do so, potentially resulting in improved treatment outcomes.

## Materials and Methods

### BC-PDOX mice

Animal experiments were approved by the West Virginia University Institutional Animal Care and Use Committee. BC-PDOX mice were produced by implanting freshly collected breast tumor tissue into the cleared mammary fat pad of NOD.CG-*Prkd*^*scid*^ *Il2rg*^*tm1 Wjl*^*/*SzJ/ 0557 (NSG) mice as previously described ^19,20^. Briefly, female NSG mice were anesthetized with isoflurane and immobilized prior to surgery. An incision was made in the skin over the lower lateral abdominal wall and a pocket was opened under the skin to expose the mammary fat pads. A 3 mm long pocket was created in the mammary fat pad, and a single tumor fragment of 2 mm^3^ was placed into the pocket. Mice were housed at 22°C under a 12-hour light/12-hour dark cycle and received food and water *ad libitum*. Each BC-PDOX model produced was authenticated with the original patient’s biopsy using genomic DNA and short tandem repeat-based PCR amplification (Arizona University Genomics Core Facility, Tucson, AZ, USA). Routine human and mouse pathogen screening was performed on original and passaged tissue by Charles River Laboratories (Wilmington, MA, USA). RNA sequence and whole exome analysis of original biopsy and passage 1 BC-PDOX further confirm relatedness of samples and stability of genomic alterations identified in BC patient and corresponding PDOX. Expression of clinically relevant BC receptors correlate with the original BC biopsy and are routinely tested by WVU Genetic and Tumor Models (GTM) Core Facility staff.

### Pioglitazone preclinical trial with BC-PDOX mice

BC-PDOX mice implanted with Her^2^/neu^+^ patient tumors (n = 16) were randomly selected to receive either pio (30mg . kg^-1^ body weight) or vehicle supplementation daily for 14 days. Tumor volume was monitored twice per week throughout the duration of the study using calipers and ultrasound (Vevo 2100). Treatment was initiated when each mouse reached a tumor volume of at least 600 mm^3^. All supplementation was delivered via oral gavage.

### RNA isolation, sequencing, and bioinformatics

Total RNA was isolated from gastrocnemius muscles of experimental mice (n=4 BC-PDOX; n=4 BC-PDOX PIO; n=3 NSC-Con) using Trizol (ThermoFisher Scientific, Waltham, MA, USA) and established methods^21^. RNA purity was assessed using a NanoDrop spectrophotometer, with 260/280 readings of at least 2.0. RNA integrity was measured on an Agilent bioanalyzer with an RNA Nano chip. RNA samples had RNA Integrity Numbers (RIN) >9, indicating high-quality RNA. RNA-Seq libraries were built using the Stranded mRNA kit from KAPA Biosciences with Illumina compatible adapters. The concentrations of the completed libraries were quantified with a qubit fluorometer using high-sensitivity DNA reagent. Libraries were subsequently run on the bioanalyzer using a high-sensitivity DNA chip. Completed libraries were then pooled in equimolar concentrations and sequenced on 1 lane of the HiSeq 1500 with PE50 bp reads and at least 20 million reads per sample. MultiQC^22^ was utilized for assessment of sequence quality and was found to be high for all sequenced samples. Salmon was used for quantification of transcript-level reads, with both gcBias and seqBias set, and libType A^23^. Differential expression analysis was conducted using DESeq2^24^ after summarizing reads to gene-level using tximport^25^. DESeq2 was run with default parameters with the exception of a filtering step: genes that did not have at least 3 samples with a count of at least 1 count per million were eliminated. RNA-Seq output was contextualized using Enrichr^26^ and The Broad Institute’s Gene Set Enrichment Analysis^27^ (GSEA) tool. Due to an RNA-processing error just prior to sequencing, final analyses were performed in n=4 BC-PDOX, n=3 BC-PDOX pio, and n=3 NSG-Con.

#### Enrichment analyses

Enrichment analyses were conducted using The Broad Institute’s GSEA tool^27^ and Enrichr^26^. Data presented in this article reflect analyses conducted in Enrichr between December 20^th^ 2020 and June 7^th^ 2021, and in GSEA in December 2020. Input data for GSEA included normalized gene expression counts for all genes with at least 1 count per million in at least 3 samples, with ranking conducted within GSEA using “Signal-to-noise” based ranking. Reference gene sets included all KEGG v7 pathways^28^ with at least 15 genes in the pathway in our gene set. Input data for Enrichr analyses included only the list of significantly differentially-expressed genes (adjusted p < 0.05). In Enrichr, all databases were queried per default settings, with particular interest in ChEA 2016 for transcription factor target enrichment and 2018 databases from the Gene Ontology project for biological process enrichment analyses. Regarding the “Combined Score” metric reported as a measure of enrichment, this value is calculated as described previously ^29^. Briefly, enrichment for a given pathway is first calculated using the Fisher exact test for many random gene lists to calculate a mean rank list for all genes. Our list of differentially-expressed genes is then compared to the expected rank, and mean and standard deviation for the difference between actual and expected rank is calculated. Combined score can then be calculated using the formula c = log(p)·z, where c is the combined score, p is the p-value computed using the Fisher exact test, and z is the z-score computed by assessing the deviation of each gene in our list of differentially-expressed genes from their corresponding expected rank. Transcription factor analysis from Enrichr is presented using the ggpubr package^30^ in R^31^.

#### Hierarchical clustering

Hierarchical clustering analysis was conducted by first calculating the Euclidean residual distance matrix for filtered, normalized, and log-transformed gene expression counts, then passing the resulting distance matrix to the heatmap.2 function of the gplots R package for dendrogram generation using the complete linkage method^32^.

#### K-means clustering

K-means unsupervised clustering analysis (k = 3, Hartigan-Wong) was conducted on the matrix of filtered, normalized, and log-transformed gene expression counts and visualized using the fviz_cluster function of the factoextra R package^33^.

### qRT-PCR validation

Total RNA was isolated from gastrocnemius muscles of BC-PDOX (n = 8) and BC-PDOX PIO (n = 8) mice. 500 ng of cDNA was produced using Invitrogen SuperScript III First-Strand Synthesis System (ThermoFisher Scientific) according to manufacturer’s protocol. Relative expression of selected genes was analyzed using SYBR Green PCR Master Mix (ThermoFisher Scientific) with the Applied Biosystems 7500 Real-Time PCR System (ThermoFisher Scientific). qRT-PCR primers **(Table 1)** were designed using Primer3^34^.

**Table 1:**
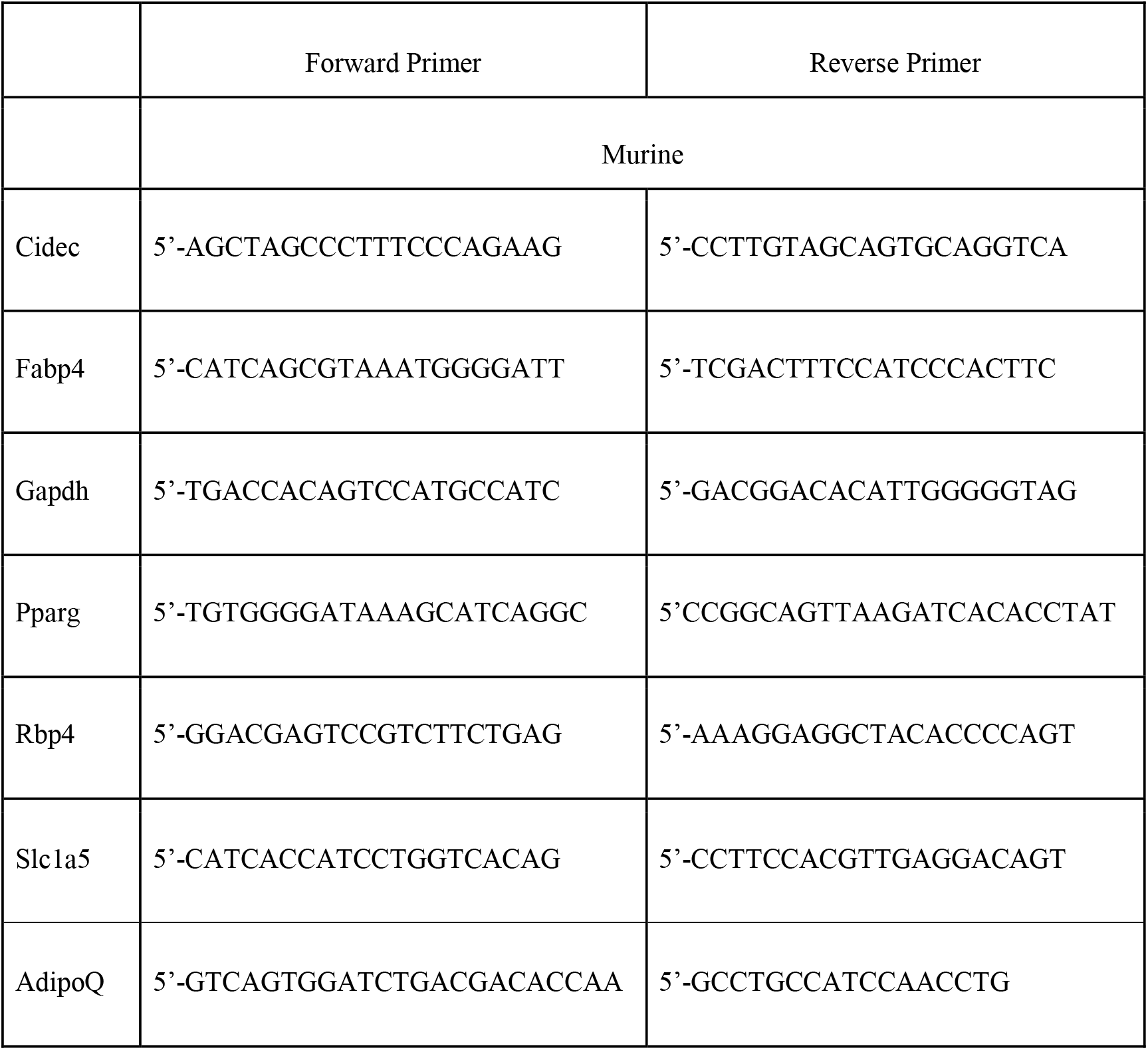
qPCR primers.

### Mitochondrial isolation and ATP analysis

Quadriceps muscles from each mouse (n = 8 BC-PDOX; n = 8 BC-PDOX PIO) were quickly removed upon euthanasia, and mitochondria were isolated using the Mammalian Mitochondria Isolation Kit for Tissue & Cultured Cells (BioVision, Cat. No. K288) according to the manufacturer’s protocol and using previously described methods^14,15,35–37^. Mitochondrial pellets were snap frozen and stored at -80C until utilized in ATP quantification and ATP synthase activity assays. ATP synthase activity was measured in isolated mitochondria, using an assay coupled with pyruvate kinase, which converts the ADP to ATP and produces pyruvate from phosphoenolpyruvate. Pyruvate produced by pyruvate kinase is converted to lactate by lactate dehydrogenase with the oxidation of NADH which is monitored at 340 nm^38^. Absorbance was measured on a Biotek Synergy HT plate reader (Biotek, Winooski, VT). The full protocol is included in the supplementary data with additional references included here^36,39^. Mitochondria ATP content was quantified using the ENLITEN® ATP Assay System Bioluminescence Detection Kit (Promega, Wisconsin, USA, using the manufacturer’s recommended standard curve for calculation of absolute ATP content from luminescence intensity and The FlexStation® 3 Multi-Mode Microplate Reader (Molecular Devices®, CA, USA). Luminescence values, reflecting ATP content, were normalized to the sample protein concentration, quantified using the DC™ Protein Assay (Bio-Rad, CA, USA).

### Causal Mediation Analysis (CMA)

A prior cohort of BC-PDOX mice were used for CMA; complete data for skeletal muscle responses in this BC-PDOX cohort can be found in^13^. Model-based CMA requires the fitting of two separate linear regression (LR) models to the data: a “mediator model,” where the mediator variable (muscle mass) is regressed on the intervention variable (tumor), and an “outcome model,” where the outcome variable (muscle fatigue, presented as AUC) is regressed on the intervention variable while controlling for the mediator. All data values for the calculation of CMA, including the area under the fatigue curve and muscle mass, are presented in **Supplemental Table 1**. Additional covariates can be added to the LR models to control for their effect on the mediator and/or the outcome variable. These LR models were then passed to the mediate function of the mediation R-package for calculation of average causal mediation effect (ACME) and average direct effect (ADE) and their associated uncertainty estimates over a number of simulations. In this analysis, 10,000 simulations per CMA were utilized for calculation of quasi-Bayesian 95% confidence intervals. Standard multiple regression was conducted in R v3.5.0^40^. Two R packages, DescTools v0.99.28^41^ and olsrr v0.5.2^42^, were utilized for appropriate assumption checking after fitting linear models. Assumptions tested include the normality of dependent variables, absence of univariate outliers, non-multicollinearity, homoscedasticity, and absence of regression outliers. No assumptions were found to be violated. Models tested are summarized as follows:

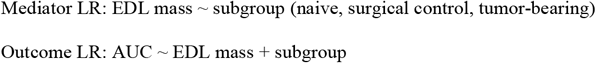

### Statistics

GraphPad Prism (V5; RRID: SCR_002798) was utilized to analyze the following data sets (qPCR, ATP quantification and ATP synthase activity, and animal weights and tumor volumes). Unless otherwise stated, all statistical tests used were unpaired, two-tailed *t* tests for two-group comparisons. When multiple comparisons were used within an experiment, Holm–Bonferroni correction was applied to adjust *P* values. Graphs represent data from a single, independent experiment. All experiments have been repeated at least twice with similar results. Data are presented as means ± SEM unless otherwise noted; statistical significance was defined a p<0.05.

## Results

### Skeletal muscle gene expression profiles in pio treated BC-PDOX mice

We have consistently observed the downregulation of a gene network in skeletal muscles from human BC patients and mice implanted with human tumor xenografts, with the transcription factor PPARG being central to this downregulated network^13–15^. To specifically target this gene network in muscle, we administered a PPARG agonist (pio) daily for 14 days in mice implanted with human Her^2^/neu^+^ tumor fragments, starting at a tumor volume of 600mm^3^ **(Fig. 1A)**. Following daily oral pio or vehicle administration, gastrocnemius muscles were processed for RNA sequencing in tumor-bearing animals and tumor-naïve NSG controls (n=4 BC-PDOX; n=3 BC-PDOX PIO, n=3 NSG-Con). Unsupervised hierarchical clustering analysis of filtered, normalized, and log-transformed RNA-seq data demonstrated that muscular gene expression patterns in 2 of the 3 BC-PDOX PIO mice clustered with non-tumor-bearing NSG-Con mice. A separate cluster was observed that included 1 of the 3 BC-PDOX PIO mice with the vehicle treated BC-PDOX mice **(Fig. 1B)**. Furthermore, principal component analysis (PCA) with k-means clustering supported these observations, with 2 of the 3 BC-PDOX PIO mice clustering with non-tumor bearing NSG-Con and separated from the vehicle-treated BC-PDOX mice (**Fig. 1C**). This analysis suggest that 2-weeks of pio-supplementation in Her^2^/neu_+_ BC-PDOX mice reverses the BC-tumor associated alterations in muscle gene expression, such that this muscle gene expression is more similar to muscle from non-tumor-bearing mice.

**Figure 1:**
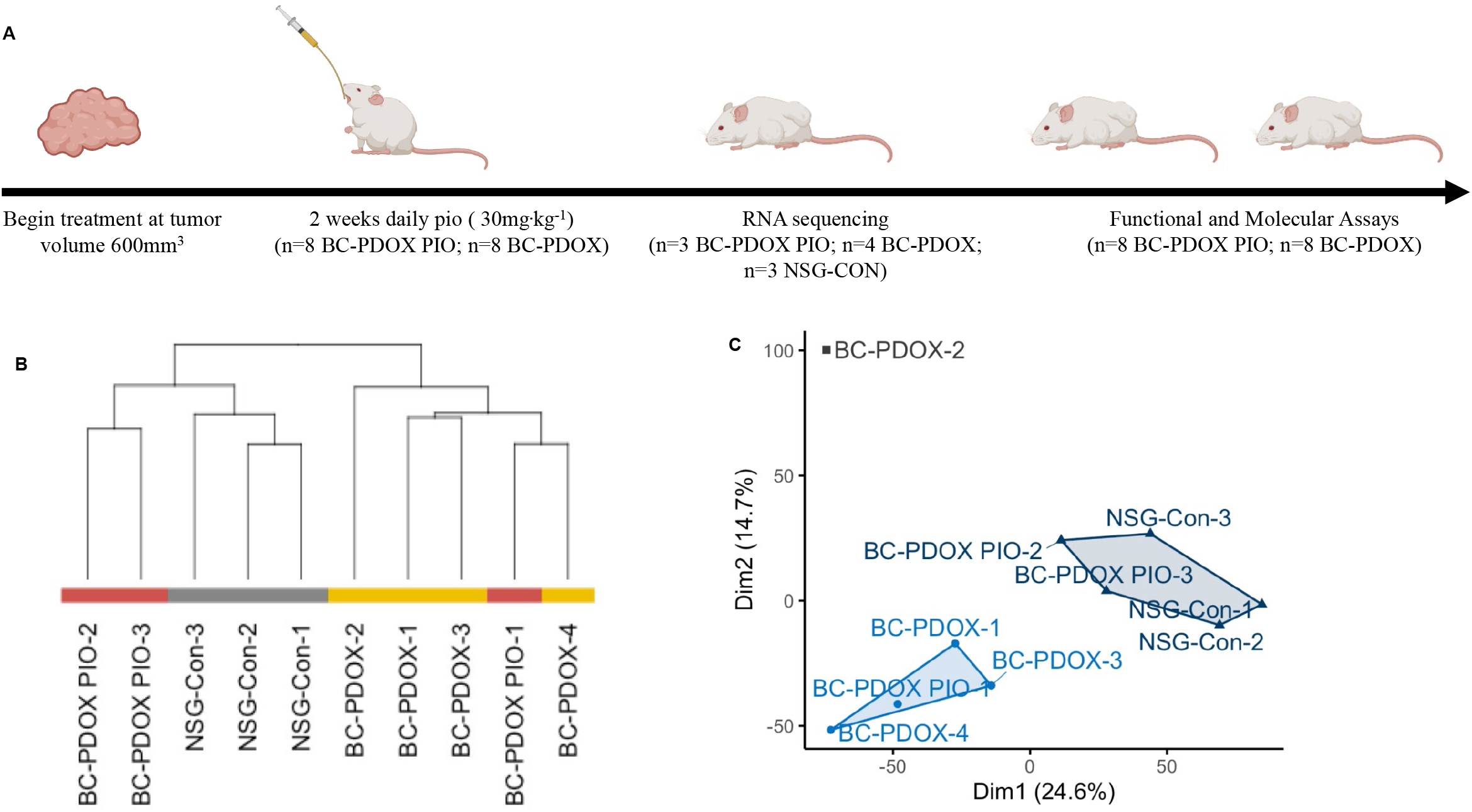
Skeletal muscle gene expression profiles in pio treated BC-PDOX mice. BC-PDOX mice were generated by implanting NSG-Con mice with human Her^2^/neu^+^ tumor fragments. Mice were randomly divided into two groups, with one group receiving supplementation of oral pio administered for 14 days (30mg/kg^-1^/day), and the second group receiving vehicle. Treatment was initiated once a tumor volume of 600mm^3^ was recorded, and all mice received oral gavage for a period of 4 weeks (**A**). Following 4 weeks of treatment, gastrocnemius muscles from BC-PDOX PIO (n=3) and BC-PDOX (n=4) treated mice were processed for RNA sequencing and compared to gastrocnemius muscle from tumor-naïve animals (n=3 NSG-Con). Unsupervised hierarchical clustering was conducted on normalized, filtered, and log-transformed RNA-seq gene expression data in muscle from study mice, with treatment group denoted by color (**B**). Principal component analysis was conducted on normalized, filtered, and log-transformed RNA-seq gene expression data in muscle from study mice with k-means (k = 3, Hartigan-Wong) clustering, with clusters denoted by point color (**C**).

### Pioglitazone reverses skeletal muscle transcriptional profile in BC-PDOX mice

Differentially-expressed genes (DEGs) in skeletal muscle were identified by comparing BC-PDOX PIO mice to non-tumor bearing NSG-Con, as well as BC-PDOX mice to non-tumor NSG-Con. A total of 592 DEGs were significantly different in muscles from BC-PDOX mice compared to muscles from non-tumor NSG-Con mice (adjusted p < 0.05) **(Supplementary Table 2)**. Conversely, a total of 174 DEGs were identified as significantly different in muscles from BC-PDOX PIO mice compared to muscles from non-tumor NSG-Con mice (adjusted p < 0.05) **(Supplementary Table 3)**. Enrichment analysis for known targets of transcription factors identified significant enrichment of genes targeted by PPARG, estrogen related receptor beta (ESRRB), interferon regulatory factor 8 (IRF8), signal transducer and activator of transcription 3 (STAT3), nuclear casein kinase and cyclin-dependent kinase substrate 1 (NUCKS1), nuclear factor E2-related factor 2 (NRF2), and liver X receptor (LXR) in muscles from BC-PDOX mice. Conversely, in BC-PDOX PIO mice, significant enrichment of genes targeted by NUCKS1 and BTB and CNC homology 1 (BACH1) was identified. Importantly, genes targeted by PPARG were not enriched in muscles from BC-PDOX PIO mice **(Figure 2A)**. These data support our prior publications and demonstrate that growth of human tumor fragments in the BC-PDOX model is associated with downregulation of a gene network regulated by PPARG^13,14^. To further explore the effects of pio administration in our model, muscle gene expression associated with biological processes was analyzed using Gene Ontology. A total of 58 dysregulated biological processes were identified in muscles from BC-PDOX mice compared to NSG-Con, while only 17 dysregulated biological processes were identified in muscles from BC-PDOX PIO mice. In muscles from BC-PDOX mice, a strong signal for aberrant mitochondrial function and electron transport chain (ETC) function is evident in the list of dysregulated biological processes (**Table 2**); these processes are not identified in muscles from BC-PDOX PIO mice (**Table 2**). Gene Set Enrichment Analysis similarly showed significant dysregulation of genes associated with oxidative phosphorylation in muscles from BC-PDOX mice and reversal of this observation in muscles of BC-PDOX PIO mice (**Fig. 2B**). Collectively, our data suggest that 14 days of oral pio therapy in BC-PDOX mice restored the muscle expression patterns to patterns more similar to those observed in the muscles of non-tumor bearing NSG-Con mice.

**Table 2:**
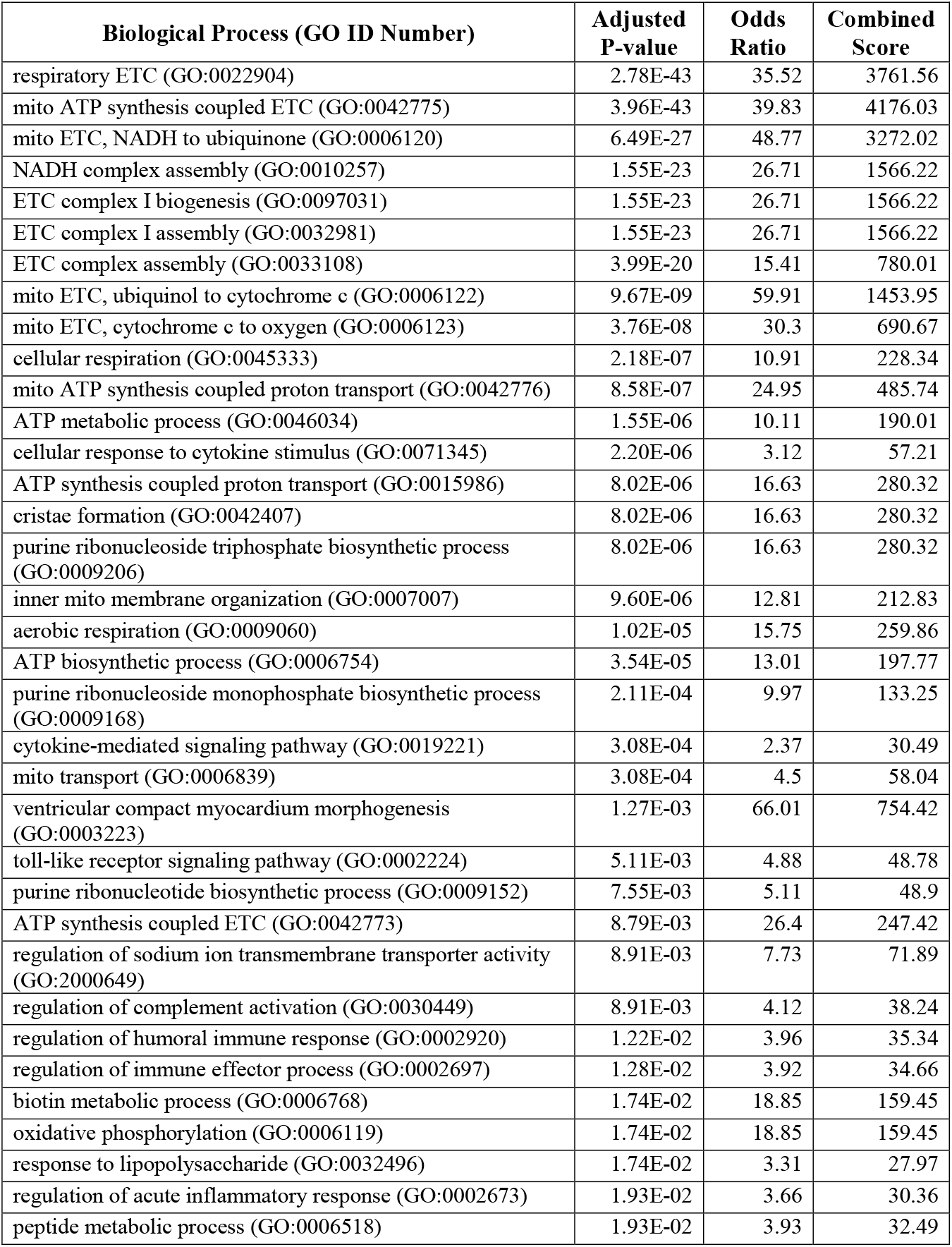
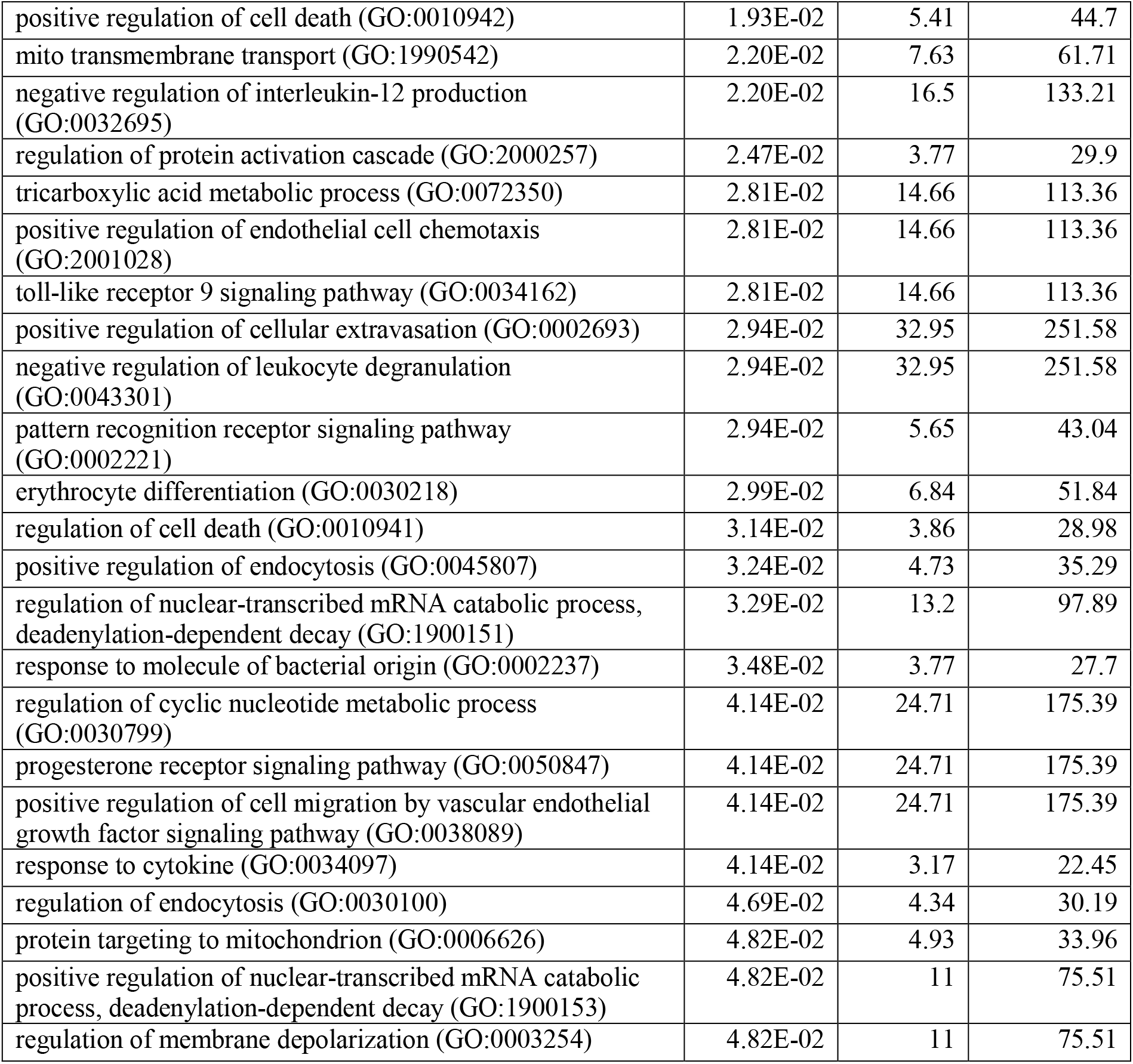
Gene Ontology Identified Biological Processes in BC-PDOX Mice.

| **Biological Process (GO ID Number)** | **Adjusted P-value** | **Odds Ratio** | **Combined Score** |
| --- | --- | --- | --- |
| respiratory ETC (GO:0022904) | 2.78E-43 | 35.52 | 3761.56 |
| mito ATP synthesis coupled ETC (GO:0042775) | 3.96E-43 | 39.83 | 4176.03 |
| mito ETC, NADH to ubiquinone (GO:0006120) | 6.49E-27 | 48.77 | 3272.02 |
| NADH complex assembly (GO:0010257) | 1.55E-23 | 26.71 | 1566.22 |
| ETC complex I biogenesis (GO:0097031) | 1.55E-23 | 26.71 | 1566.22 |
| ETC complex I assembly (GO:0032981) | 1.55E-23 | 26.71 | 1566.22 |
| ETC complex assembly (GO:0033108) | 3.99E-20 | 15.41 | 780.01 |
| mito ETC, ubiquinol to cytochrome c (GO:0006122) | 9.67E-09 | 59.91 | 1453.95 |
| mito ETC, cytochrome c to oxygen (GO:0006123) | 3.76E-08 | 30.3 | 690.67 |
| cellular respiration (GO:0045333) | 2.18E-07 | 10.91 | 228.34 |
| mito ATP synthesis coupled proton transport (GO:0042776) | 8.58E-07 | 24.95 | 485.74 |
| ATP metabolic process (GO:0046034) | 1.55E-06 | 10.11 | 190.01 |
| cellular response to cytokine stimulus (GO:0071345) | 2.20E-06 | 3.12 | 57.21 |
| ATP synthesis coupled proton transport (GO:0015986) | 8.02E-06 | 16.63 | 280.32 |
| cristae formation (GO:0042407) | 8.02E-06 | 16.63 | 280.32 |
| purine ribonucleoside triphosphate biosynthetic process (GO:0009206) | 8.02E-06 | 16.63 | 280.32 |
| inner mito membrane organization (GO:0007007) | 9.60E-06 | 12.81 | 212.83 |
| aerobic respiration (GO:0009060) | 1.02E-05 | 15.75 | 259.86 |
| ATP biosynthetic process (GO:0006754) | 3.54E-05 | 13.01 | 197.77 |
| purine ribonucleoside monophosphate biosynthetic process (GO:0009168) | 2.11E-04 | 9.97 | 133.25 |
| cytokine-mediated signaling pathway (GO:0019221) | 3.08E-04 | 2.37 | 30.49 |
| mito transport (GO:0006839) | 3.08E-04 | 4.5 | 58.04 |
| ventricular compact myocardium morphogenesis (GO:0003223) | 1.27E-03 | 66.01 | 754.42 |
| toll-like receptor signaling pathway (GO:0002224) | 5.11E-03 | 4.88 | 48.78 |
| purine ribonucleotide biosynthetic process (GO:0009152) | 7.55E-03 | 5.11 | 48.9 |
| ATP synthesis coupled ETC (GO:0042773) | 8.79E-03 | 26.4 | 247.42 |
| regulation of sodium ion transmembrane transporter activity (GO:2000649) | 8.91E-03 | 7.73 | 71.89 |
| regulation of complement activation (GO:0030449) | 8.91E-03 | 4.12 | 38.24 |
| regulation of humoral immune response (GO:0002920) | 1.22E-02 | 3.96 | 35.34 |
| regulation of immune effector process (GO:0002697) | 1.28E-02 | 3.92 | 34.66 |
| biotin metabolic process (GO:0006768) | 1.74E-02 | 18.85 | 159.45 |
| oxidative phosphorylation (GO:0006119) | 1.74E-02 | 18.85 | 159.45 |
| response to lipopolysaccharide (GO:0032496) | 1.74E-02 | 3.31 | 27.97 |
| regulation of acute inflammatory response (GO:0002673) | 1.93E-02 | 3.66 | 30.36 |
| peptide metabolic process (GO:0006518) | 1.93E-02 | 3.93 | 32.49 |
| positive regulation of cell death (GO:0010942) | 1.93E-02 | 5.41 | 44.7 |
| mito transmembrane transport (GO:1990542) | 2.20E-02 | 7.63 | 61.71 |
| negative regulation of interleukin-12 production (GO:0032695) | 2.20E-02 | 16.5 | 133.21 |
| regulation of protein activation cascade (GO:2000257) | 2.47E-02 | 3.77 | 29.9 |
| tricarboxylic acid metabolic process (GO:0072350) | 2.81E-02 | 14.66 | 113.36 |
| positive regulation of endothelial cell chemotaxis (GO:2001028) | 2.81E-02 | 14.66 | 113.36 |
| toll-like receptor 9 signaling pathway (GO:0034162) | 2.81E-02 | 14.66 | 113.36 |
| positive regulation of cellular extravasation (GO:0002693) | 2.94E-02 | 32.95 | 251.58 |
| negative regulation of leukocyte degranulation (GO:0043301) | 2.94E-02 | 32.95 | 251.58 |
| pattern recognition receptor signaling pathway (GO:0002221) | 2.94E-02 | 5.65 | 43.04 |
| erythrocyte differentiation (GO:0030218) | 2.99E-02 | 6.84 | 51.84 |
| regulation of cell death (GO:0010941) | 3.14E-02 | 3.86 | 28.98 |
| positive regulation of endocytosis (GO:0045807) | 3.24E-02 | 4.73 | 35.29 |
| regulation of nuclear-transcribed mRNA catabolic process, deadenylation-dependent decay (GO:1900151) | 3.29E-02 | 13.2 | 97.89 |
| response to molecule of bacterial origin (GO:0002237) | 3.48E-02 | 3.77 | 27.7 |
| regulation of cyclic nucleotide metabolic process (GO:0030799) | 4.14E-02 | 24.71 | 175.39 |
| progesterone receptor signaling pathway (GO:0050847) | 4.14E-02 | 24.71 | 175.39 |
| positive regulation of cell migration by vascular endothelial growth factor signaling pathway (GO:0038089) | 4.14E-02 | 24.71 | 175.39 |
| response to cytokine (GO:0034097) | 4.14E-02 | 3.17 | 22.45 |
| regulation of endocytosis (GO:0030100) | 4.69E-02 | 4.34 | 30.19 |
| protein targeting to mitochondrion (GO:0006626) | 4.82E-02 | 4.93 | 33.96 |
| positive regulation of nuclear-transcribed mRNA catabolic process, deadenylation-dependent decay (GO:1900153) | 4.82E-02 | 11 | 75.51 |
| regulation of membrane depolarization (GO:0003254) | 4.82E-02 | 11 | 75.51 |

**Table 3:** Gene Ontology Identified Biological Processes in BC-PDOX Mice Treated with Pioglitazone.

| **Biological Process (GO ID Number)** | **Adjusted P-value** | **Odds Ratio** | **Combined Score** |
| --- | --- | --- | --- |
| phospholipid transport (GO:0015914) | 5.52E-04 | 16.25 | 230.53 |
| phospholipid translocation (GO:0045332) | 5.52E-04 | 36.63 | 511.8 |
| lipid translocation (GO:0034204) | 5.52E-04 | 34.47 | 473.03 |
| cellular response to cytokine stimulus (GO:0071345) | 9.48E-04 | 4.46 | 57.53 |
| response to lipid (GO:0033993) | 9.49E-03 | 7.19 | 74.54 |
| response to lipopolysaccharide (GO:0032496) | 1.63E-02 | 6.45 | 62.21 |
| regulation of cholesterol storage (GO:0010885) | 2.57E-02 | 38.63 | 344.37 |
| cytokine-mediated signaling pathway (GO:0019221) | 2.57E-02 | 3.15 | 28.07 |
| MAPK cascade (GO:0000165) | 2.62E-02 | 4.45 | 38.7 |
| positive regulation of cardiac muscle cell proliferation (GO:0060045) | 2.62E-02 | 34.76 | 301.02 |
| response to cytokine (GO:0034097) | 2.76E-02 | 6.3 | 53.42 |
| interleukin-6-mediated signaling pathway (GO:0070102) | 2.76E-02 | 31.6 | 266.22 |
| JAK-STAT cascade involved in growth hormone signaling pathway (GO:0060397) | 3.16E-02 | 28.97 | 237.75 |
| positive regulation of cardiac muscle tissue growth (GO:0055023) | 3.59E-02 | 26.74 | 214.07 |
| reverse cholesterol transport (GO:0043691) | 4.04E-02 | 24.83 | 194.1 |
| regulation of cardiac muscle cell proliferation (GO:0060043) | 4.52E-02 | 23.17 | 177.08 |
| positive regulation of peptidyl-tyrosine phosphorylation (GO:0050731) | 4.75E-02 | 6.4 | 48.22 |

**Figure 2:**
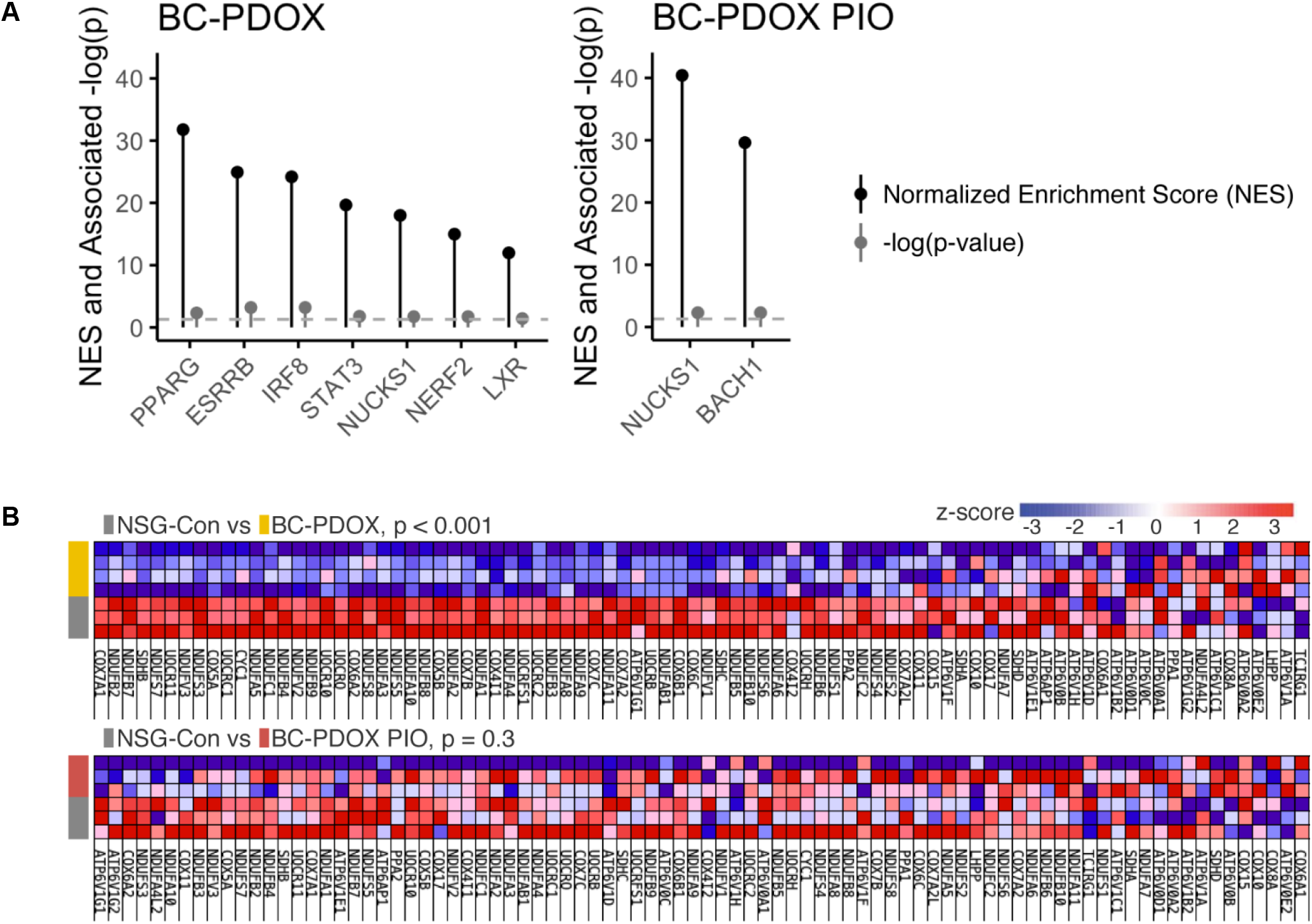
Pioglitazone reverses skeletal muscle transcriptional changes in BC-PDOX mice. Differentially-expressed genes (DEGs) in skeletal muscle were identified. Significantly enriched (adjusted p < 0.05) transcription factors in BC-PDOX group vs NSG-Con (left) and in BC-PDOX PIO group vs NSG-Con (right), querying ChEA 2016 transcription factor gene sets via Enrichr (**A**). Heatmap representing gene expression z-scores of genes within KEGG Oxidative Phosphorylation pathway (hsa00190) in individual mice from BC-PDOX vs NSG-Con (top) and BC-PDOX PIO vs NSG-Con (bottom), queried via GSEA, with treatment group denoted by color. Individual genes are ordered from left to right according to “signal-to-noise” based ranking per GSEA’s algorithm (**B**).

### Validation of RNA-Seq profiles in the skeletal muscle of BC-PDOX mice

The muscle gene expression of PPARG and several verified PPARG target genes were quantified by qRT-PCR in muscles from BC-PDOX PIO mice (n=7) and BC-PDOX mice (n=8) to validate the RNA-Seq analyses. The muscle samples utilized in this validation were a combination of muscles used in RNA-Seq experiments and independent muscle samples from BC-PDOX mice. Pio supplementation in BC-PDOX PIO significantly increased the muscle expression of all PPARG target genes (*Cidec, Fabp4, Rbp4, Slc1a5, AdipoQ)* compared to muscles from BC-PDOX vehicle-treated mice (**Fig. 3A-E**). The expression of PPARG was not significantly changed by pio treatment, consistent with our previous work^13^ and the mechanism of action of pio **(Fig 3F)**. These observations provide support for the use of pio to reverse the downregulation of this PPARG-associated gene network in skeletal muscle in BC models. Furthermore, as we have consistently observed downregulation of these PPARG target genes in skeletal muscles of human BC-patients and in the BC-PDOX model^13,14,43^, we propose that this gene subset may be considered a “biomarker” for BC-associated fatigue in patients with early stage BC.

**Figure 3:**
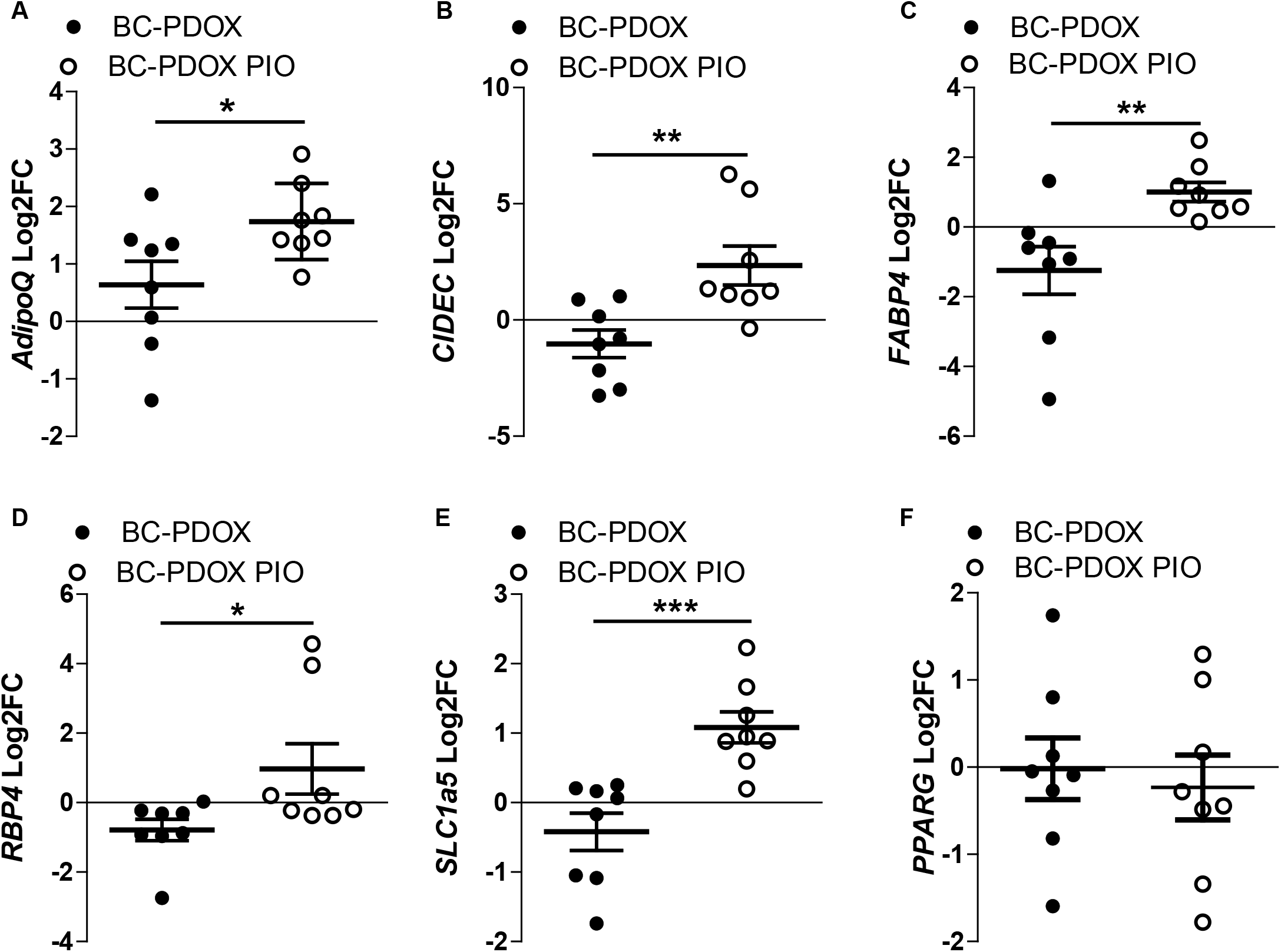
Validation of RNA-Seq profiles in the skeletal muscle of BC-PDOX mice. The gene expression of PPARG and several previously verified PPARG target genes were validated by qRT-PCR in the same samples used in RNA-Seq (n = 4 BC-PDOX PIO, n = 4 BC-PDOX) as well as in skeletal muscle from a new cohort of BC-PDOX mice (n = 4 BC-PDOX PIO, n = 4 BC-PDOX). PPARG target genes expression (*AdipoQ, Cidec, Fabp4, Rbp4, Slc1a5,PPARG)* compared to vehicle-treated mice (**A-F**) in both cohorts. 3 technical replicates of each sample were used for each gene assayed, and results were replicated across 2 individual plates. Adjusted p-values represent Holm-Bonferroni adjusted p-values of paired t-tests, n = 8 per group, all values are means ± SEM. * p < 0.05, ** p < 0.01, *** p <0.001.

### Pioglitazone supplementation restores skeletal muscle ATP content and ATP synthase activity

Live mitochondria were isolated from the quadriceps muscle of BC-PDOX PIO (n = 8) and BC-PDOX (n = 8) mice and were processed for the quantification of ATP content as well as evaluation of ATP synthase enzyme activity. We found that skeletal muscle mitochondria from BC-PDOX PIO mice had a greater ATP content compared to vehicle treated BC-PDOX mice (**Fig. 4A)**. Similarly, the activity of ATP synthase was significantly greater in mitochondria from BC-PDOX PIO mice compared to mitochondria from BC-PDOX mice (**Fig. 4B**), an observation consistent with the increased overall ATP content. These data indicate that skeletal muscle ATP and ATP synthase activity may be directly related to increased expression of PPARG target genes in skeletal muscle. Furthermore, these data suggest that pio supplementation may benefit patients with BC-associated muscle fatigue by improving ATP availability within skeletal muscle.

**Figure 4:**
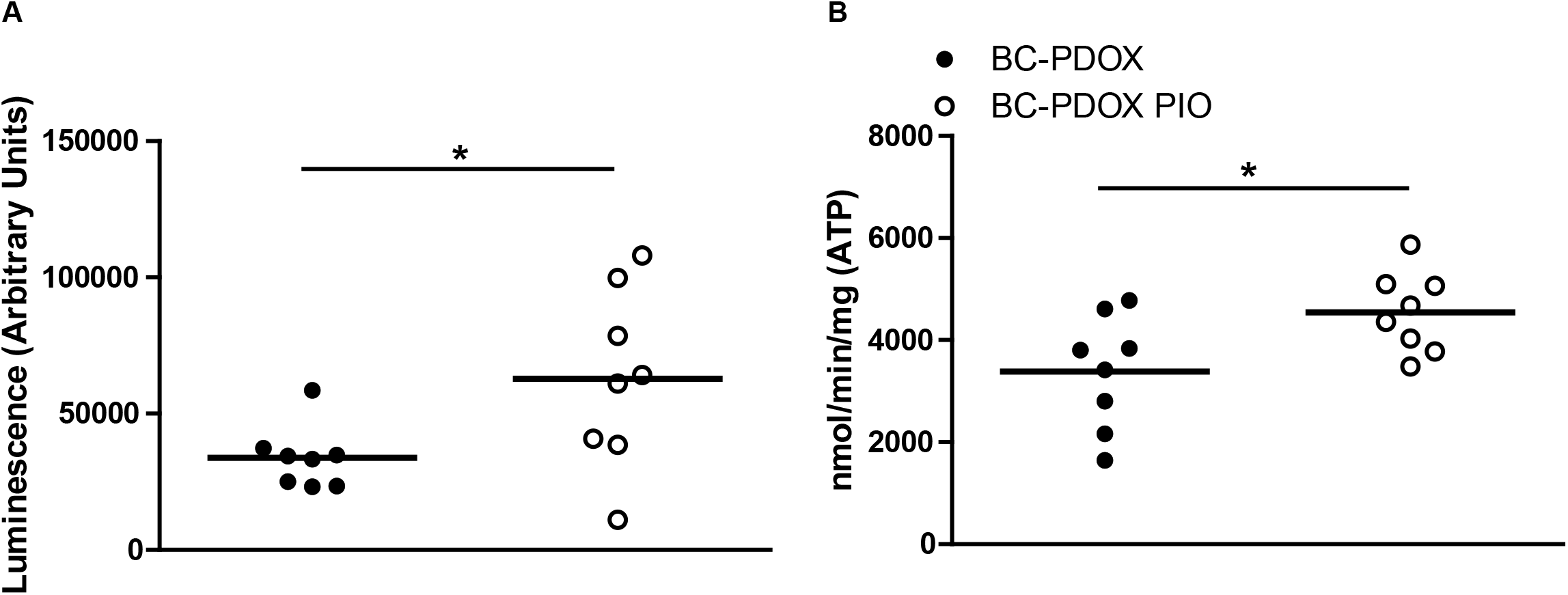
Pioglitazone supplementation restores skeletal muscle ATP and ATP synthase activity. Live mitochondria were isolated form the quadriceps muscle of BC-PDOX PIO (n = 8) and BC-PDOX (n = 8) treated BC-PDOX mice. Luminescence intensity was calculated for the quantification of ATP content (**A**). ATP synthase activity was measured (**B**) using a spectrophotometric assay. Absorbance was measured on a Biotek Synergy HT plate reader (Biotek, Winooski, VT) ETC Complex V Specific Activity (nmol min^-1^ mg^-1^) = (Δ Absorbance/min x 1,000) / [(extinction coefficient x volume of sample used in ml) x (sample protein concentration in mg ml^-1^)] where extinction coefficient for NADH is 6.2 mM^-1^ cm^-1^. ATP content and ATP synthase activity in the BC-PDOX PIO treated mitochondria was compared to BC-PDOX controls using the Mann-Whitney U test, n = 8 per group, all values are means ± SEM. * p < 0.05.

### Characteristics of BC-PDOX mice following pio supplementation

Tumor growth progressed linearly during the 2-weeks of pio supplementation, with no differences observed in BC-PDOX mice compared to BC-PDOX PIO mice **(Fig 5A)**. Body weight remained consistent during the 2-weeks of pio supplementation, with no differences between BC-PDOX mice and BC-PDOX PIO mice **(Fig 5B)**. At the time of euthanasia, no differences were observed in the tumor weight of BC-PDOX mice compared to BC-PDOX PIO mice **(Fig 5C)**. Similarly, no differences were observed in the tumor-free body weight of BC-PDOX mice compared to BC-PDOX PIO mice at euthanasia **(Fig 5D)**. Skeletal muscle mass for selected hindlimb muscles, including the gastrocnemius, solues, tibialis anterior, and extensor digitorum longus, were not different when comparing BC-PDOX mice to BC-PDOX PIO mice at euthanasia. Collectively, these data support our prior observations, that human breast tumor growth in the BC-PDOX model is not associated with significant body weight loss or muscle wasting. Furthermore, these data suggest that 2 weeks of daily pio supplementation does not affect tumor growth in the BC-PDOX model.

**Figure 5:**
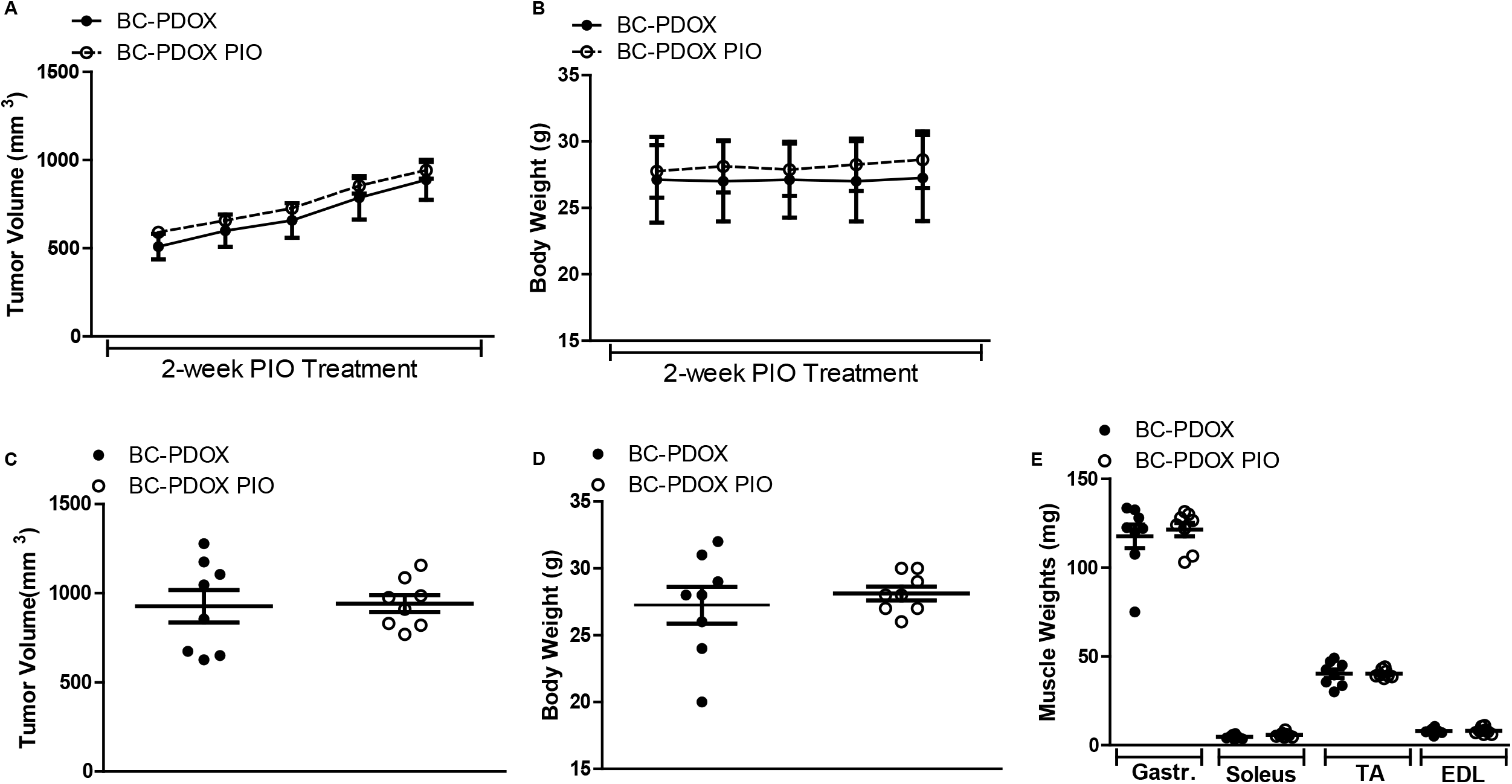
Characteristics of BC-PDOX mice following pio supplementation. Tumor volumes and body weights of BC-PDOX mice were tracked throughout the study, and muscle weights were obtained upon euthanasia. The rates of tumor growth (**A**) and body weight change (**B**), final tumor volumes (**C**) final body weights (**D**), and individual muscle weights (**E**) were recorded. Adjusted p-values represent Holm-Bonferroni adjusted p-values of paired t-tests, n = 8 per group, all values are means ± SEM. Linear regression was performed to calculate the slopes in (**A and B**).

### Skeletal muscle fatigue in BC-PDOX mice is not a function of cachexia

A consistent observation of the BC-PDOX model has been a greater rate of muscle fatigue without concomitant muscle wasting^13–15^. This phenotype resembles observations in patients with BC^13^ but differs from other murine models in which muscle fatigue is documented with associated muscle wasting^4,44,45^ To statistically evaluate the contribution of changes in muscle mass to muscle fatigue in our model, causal mediation analysis (CMA) was performed in a previous cohort of muscles from BC-PDOX mice (n = 6) using *ex vivo* muscle fatigue methodology^13^ (**Fig. 6A**). Muscles from naive NSG mice (n = 7) and mice that were implanted with BC-PDOXs that did not engraft (n = 9) were used as controls for the CMA analysis. Three different models of CMA were used to calculate the average direct effects (ADE) of tumor growth on muscle fatigue and the average causal mediator effects (ACME) of muscle mass on muscle fatigue in our model. In the three different CMA models tested, there were significant values for the direct effect (ADE) of tumor growth on the muscle fatigue response, with a negative effect size in all cases, indicating that tumor growth results in a decreased area under the curve value that reflects increased fatigability **(Table 4)**. However, muscle mass was not a significant mediator (ACME) of the muscle fatigue response. The ADE was strongest when comparing the BC-PDOX mice and the naive control mice **(Fig. 6B)**, while the comparison of BC-PDOX mice to surgical controls reached a p-value equal to 0.05 **(Fig. 6C)**. When both control groups were combined into a single group, the ADE was highly significant **(Fig. 6D)**. This statistical analysis suggests that tumor growth in the BC-PDOX mouse model directly contributes to the observed greater muscle fatigue and that loss of muscle mass is likely not a mediator of muscle fatigue in this mouse model of BC.

**Table 4:** Summary of Causal Mediation Analysis Models.

| Model | Comparison | ACME<br>Estimate<br>(Muscle Mass)<br>(95% CI) | ACME<br>p-value | ADE Estimate<br>(Fatigue)<br>(95% CI) | ADE<br>p-value |
| --- | --- | --- | --- | --- | --- |
| **BC-PDOX** | All Control vs. PDOX | -150 (-655, 167) | 0.42 | -1650 (-2500, -831) | <0.0001*** |
|  | Naïve Control vs. PDOX | -147 (-664, 171) | 0.43 | -2080 (-2940, -1233) | <0.0001*** |
|  | Surgical Control vs. PDOX | -170 (-717, 156) | 0.38 | -886 (-1767, 7) | 0.05 |

**Figure 6:**
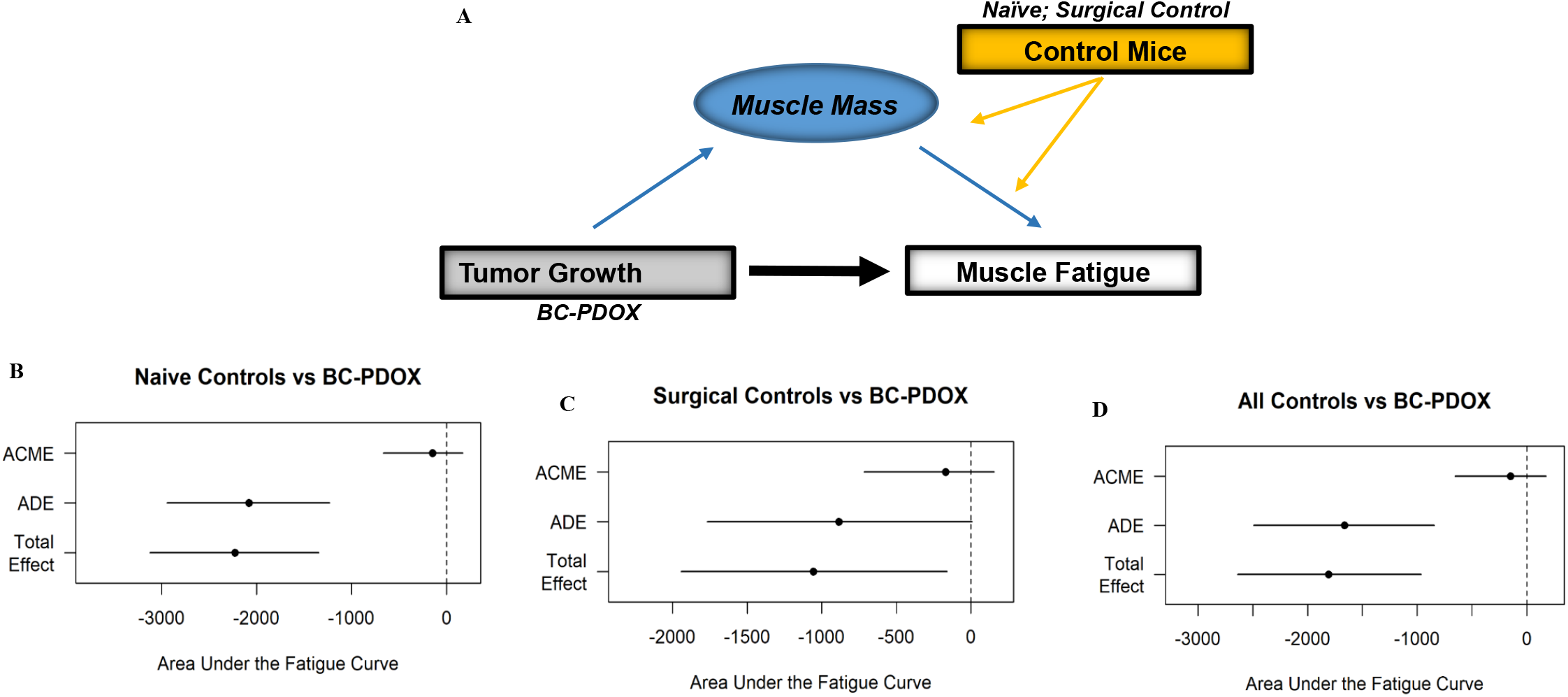
Skeletal muscle fatigue in BC-PDOX mice is not a function of cachexia. Causal mediation analysis (CMA) was performed in a previous cohort of BC-PDOX mice (n = 6) (**A**). Muscles from naïve NSG-Con mice (n = 7, naïve controls) and mice implanted with BC-PDOXs that did not engraft (n = 9, surgical controls) were used as controls for the CMA analysis. The ADE was calculated comparing the BC-PDOX mice and the naïve control mice (**B**), while the comparison of BC-PDOX mice to surgical controls reached a p-value equal to 0.05 (**C**). Both groups combined into a single group is shown in (**D**).

## Discussion

According to the latest statistics from the National Cancer Institute, approximately 13% of women will be diagnosed with BC at some point in their lives, and as of 2018, approximately 3.6 million women were living with BC in the United States^46^. Improved BC surveillance has contributed to a lower percentage of BC-patients presenting with metastatic disease at the time of diagnosis, and primary BC treatments are becoming increasingly effective, with 5-year survival rates of over 90% from 2011-2017 ^46^. While these statistics are promising, the vast majority of women diagnosed with BC experience cancer-related fatigue, and cancer-related fatigue is known to contribute to adverse outcomes by reducing a patient’s physical and psychological tolerance for anti-cancer therapies such as chemotherapy and radiation^5,6,8,10,47^. Therefore, supportive therapies specifically targeting fatigue could improve both quality of life and survival in BC. While the incidence of fatigue in BC patients is well-known, clinical diagnosis and intervention for fatigue occurs infrequently. This may reflect the assumption that cancer-related muscle fatigue is causally linked to the broader paraneoplastic syndrome known as cancer cachexia. This is of particular relevance to BC patients, as incidence of frank cachexia, defined as weight loss exceeding 5% of body weight, is quite low in BC^2,4^. Therefore, any potentially targetable mechanism underlying fatigue that is intrinsic to skeletal muscle remains unaddressed, as interventions are typically only prescribed for patients with cachexia-induced weight loss. This discrepancy highlights the need to address the unique mechanisms that contribute to BC-associated fatigue in the absence of frank cachexia, with the goal of early diagnosis and intervention aimed at improving overall treatment outcomes.

Data from our laboratory in muscle from human BC patients and the BC-PDOX mouse model have identified a BC-induced skeletal muscle phenotype that includes widespread transcriptional changes, mitochondrial dysfunction, and fatigue^13–15,43^. These studies have consistently identified downregulation of PPARG as central to this phenotype, an observation that is independent of BC molecular subtype (i.e. luminal, triple negative, Her^2^/neu^+^) and exposure to tumor-directed treatments (i.e. chemotherapy, radiation) ^14^. We hypothesized that supplementation with the PPARG agonist pio would increase the skeletal muscle expression of PPARG target genes involved in mitochondrial regulatory networks, thereby potentially improving BC-associated fatigue. In vehicle-treated BC-PDOX mice, we once again identified downregulation of gene networks regulated by PPARG, but these changes were not identified in BC-PDOX PIO mice following 2 weeks of pio supplementation. Furthermore, gene expression patterns in muscles from BC-PDOX PIO were restored to the expression patterns of muscles from non-tumor bearing NSG-Con mice. Gene ontology analysis identified 58 dysregulated biological pathways in muscles from vehicle-treated BC-PDOX mice compared to NSG-Con, while only 17 dysregulated biological processes were identified in muscles of BC-PDOX PIO mice. Among these identified biological processes, only vehicle-treated BC-PDOX mice exhibited a signal for aberrant mitochondrial function and oxidative phosphorylation, with correction of this in BC-PDOX PIO mice. Finally, we observed both increased ATP and ATP synthase activity in the mitochondria of BC-PDOX PIO mice, providing direct support for the role of pio in improving skeletal muscle mitochondrial function in our BC-PDOX model. Collectively, our data support a reversal of the BC-associated molecular alterations in skeletal muscle with short-term pio supplementation; an observation that would be consistent with an improvement in cancer-associated fatigue.

The underlying mechanisms of cancer-associated fatigue are not well characterized. The ability of tumors to induce structural, histopathological, and molecular changes in distant tissues is becoming increasingly recognized^48^, and tumor growth induces systemic metabolic changes including the development of insulin resistance^45,49^, and altered glucose and lipid metabolism^50^. The organ system that is primarily involved in the regulation of these metabolic processes is the skeletal muscle, and previous studies have described BC-induced skeletal muscle transcriptional and mitochondrial changes and associated fatigue^13,15^. The link between systemic skeletal muscle changes and localized breast tumor growth may be the function of PPARG, a nuclear transcription factor that has demonstrated roles in whole body energy regulation, regulation of insulin sensitivity, lipid metabolism and transport, and glucose metabolism^16–18^. We have consistently demonstrated that a systemic consequence of breast tumor growth is the downregulation of PPARG in skeletal muscle^13,14^, and this phenotype can also be induced by applying isolated BC-derived factors to skeletal muscle cells *in vitro*^15^. This interaction results in the aberrant transcription of muscle mitochondrial genes and concomitant mitochondrial dysfunction. Proteomics performed on the muscle of BC patients with Her2/neu^+^ tumors has demonstrated that nearly every protein involved in the mitochondrial ETC is downregulated relative to women without BC, and this is accompanied by decreased mitochondrial ATP content^14^. Additional studies have supported this mitochondrial dysfunction, as BC conditioned media applied to C2C12 myotubes has been shown to decrease mitochondrial respiration and electron flow^15^, which can be reversed by exogenous expression of PPARG. PPARG as a therapeutic target has also been shown to rescue mitochondrial dysfunction in a number of neurological disorders, including ALS^51^, and Huntington’s disease^52,53^. In a model of Alzheimer’s disease, PPARG agonist treatment increased mitochondrial biogenesis and improved glucose utilization^52^. Additionally rosiglitazone treatment increased mitochondrial biogenesis, increased oxygen consumption, increased mitochondrial mass, ΔΨm, mtDNA copy number, decreased autophagy and suppressed free radical generation in a model of Parkinson’s disease^54^. Pioglitazone also has a well-established role in the treatment of type 2 diabetes, and has been shown to increase the expression of skeletal muscle mitochondrial genes^55^, improve mitochondrial oxidative function^56^, and correct dysregulation of skeletal muscle mitochondrial proteins involved in ATP synthesis^57^. In support of the mechanisms demonstrated in these previous studies, the current study demonstrates therapeutically targeting PPARG in BC with pio is sufficient to restore muscle gene expression patterns to that observed in muscles from non-tumor bearing control mice. This is particularly evident through the restoration of the transcription of genes involved in pathways associated with oxidative phosphorylation. This transcriptional restoration is accompanied by increased mitochondrial ATP content and ATP synthase activity. Given the relationship between proper skeletal muscle metabolic function and ATP generation, to the incidence of fatigue, and the role of pio and other PPARG agonists in improving mitochondrial function in other diseases, we suggest that pio be considered a promising candidate for the amelioration of fatigue in patients with BC.

The use of biomarkers is common in many cancers, and these typically constitute a biological molecule found in blood, other body fluids, or tissues that is a sign of a normal or abnormal process^58^. Biomarkers have many potential applications in clinical oncology, including risk assessment, screening, diagnosis, determination of prognosis, prediction of response to treatment, and monitoring of progression of disease. There are a tremendous variety of biomarkers, which can include proteins, nucleic acids, antibodies, and peptides. For example, the expression of BRCA is a well-known biomarker in breast cancer^59^. We have identified a PPARG target gene panel (*FABP4, CIDEC, SLC1a5, RBP4*) that is consistently downregulated in skeletal muscle from BC patients, as well as multiple cohorts of BC-PDOX mice^13,14,43^. Further, these results have remained consistent across all breast tumor molecular subtypes, and independent of exposure to chemotherapy or radiation^14^. We have demonstrated that the altered gene expression patterns and associated mitochondrial dysfunction and muscle fatigue occurs in the absence of weight loss. Therefore, we propose that the identified PPARG target gene panel (*FABP4, CIDEC, SLC1a5, RBP4*) be considered as a diagnostic biomarker for BC-associated skeletal muscle fatigue in weight stable patients with early stage disease.

Causal mediation analysis is a statistical technique that is based on causal effects, makes certain assumptions in the modeling explicit, and may be applied to nonparametric or curvilinear data^60,61^. The statistical modeling allows one to control for the effects on a confounding mediator on the outcome variable being measured^60^. For example, a recent study used CMA to provide an unbiased estimate of the effect of muscle hypertrophy on overall muscle strength levels in response to resistance training^60^. In the current study, we utilized CMA to estimate the effects of muscle mass loss (i.e. cachexia) on the rate of muscle fatigue in response to breast tumor growth in our BC-PDOX model. Our initial statistical analysis comprised 3 different CMA models based on data acquired in two cohorts of control mice. The CMA analysis presented in this work suggests that breast cancer tumor growth contributes directly to fatigue, and that muscle mass is not likely a mediator of muscle fatigue in BC-PDOX mice. Importantly, these analyses were statistically significant for all three models tested with associated negative effect sizes, providing evidence that breast tumor growth directly contributes to the rate of muscle fatigue and that muscle mass loss is not a confounding mediator of this relationship. These data in the BC-PDOX model are reflective of the clinical observation that while cachexia is relatively rare in BC-patients, fatigue is often one of the first recognized symptoms of BC, and is often present prior to a BC diagnosis^4^.

Overall, the data presented in this study support the therapeutic potential of pio for the treatment of BC-induced muscle fatigue through the restoration of skeletal muscle gene expression patterns and mitochondrial function. A limitation of the study is that skeletal muscle fatigability was not directly measured following the pio treatment; however, this will be assessed in a future cohort of BC-PDOX mice. We expect that the increased mitochondrial ATP content and ATP synthase activity demonstrated with pio treatment in the current study will contribute to a reduction in muscle fatigue in our model. In conclusion, we have demonstrated that the PPARG agonist pio, is capable of reversing the expected gene expression patterns in the skeletal muscle of PDOX mice, as well as improving ATP availability in muscle. With this, further studies are warranted to determine whether pio can directly improve skeletal muscle fatigue and provide quality of life benefits to BC patients.

## Acknowledgements

This research was supported by the following: National Institute of General Medical Sciences of the National Institutes of Health under Award Number P20GM121322 (Lockman), American Cancer Society Institutional Research Grant 09-061-04 (Pistilli). Authors would like to acknowledge the following WVU associated facilities: Preclinical Tumor Models Core Facility (CA148671, Pugacheva); Mitochondria Core of the WVU Stroke CoBRE (P20GM109098); Mitochondria, Metabolism and Bioenergetics group (R01 HL-128485; Hollander and the Community Foundation for the Ohio Valley Whipkey Trust); Genomics Core Facility (CTSI Grant U54 GM104942).

## Figure Legends

**Table 1: BC-PDOX Dysregulated Biological Processes**

All significantly enriched biological processes (Benjamin-Hochberg adjusted p-value < 0.05) in the set of differentially expressed genes in BC-PDOX vs. NSG-Con mice, from Gene Ontology Project Biological Processes 2018 queried via Enricher. Odds ratio and associated p-value for enrichment was computed using Fisher exact test. Combined Score represents a second way of computing enrichment, described under Methods and referenced in^29^. Higher Combined Scores and higher Odds Ratios indicate a greater degree of enrichment.

**Table 2: BC-PDOX PIO Dysregulated Biological Processes**

All significantly enriched biological processes (Benjamin-Hochberg adjusted p-value < 0.05) in the set of differentially expressed genes in BC-PDOX PIO vs. NSG-Con mice, from Gene Ontology Project Biological Processes 2018 queried via Enricher. Odds ratio and associated p-value for enrichment was computed using Fisher exact test. Combined Score represents a second way of computing enrichment, described under Methods and referenced in^29^. Higher Combined Scores and higher Odds Ratios indicate a greater degree of enrichment.

