## Supplemental Table 1 for "Pioglitazone administration restores a PPARG-dependent transcriptional network and ATP levels within skeletal muscles of mice implanted with patient-derived breast tumors"

**Supplemental Table 1: Area Under the Curve values and EDL muscle mass from Control and PDX tumor-bearing mice for CMA calculations.**

| Mouse ID # | Control or Tumor | Limb Used | AUC | EDL Mass (mg) |
|---|---|---|---|---|
| NSG6 | con | R | 10099 | 14.7 |
| | con | L | 7623 | 17.6 |
| NSG7 | con | R | 9775 | 13.5 |
| NSG8 | con | R | 8289 | 8.5 |
| NSG9 | con | R | 7915 | 6.6 |
| NSG10 | con | R | 8236 | 7 |
| NSG11 | con | R | 9135 | 5.9 |
| SCon1 | Surgical Con | R | 7477 | 10 |
| SCon2 | Surgical Con | R | 7633 | 9.4 |
| SCon3 | Surgical Con | R | 7996 | 9.8 |
| SCon4 | Surgical Con | R | 6671 | 9.7 |
| | Surgical Con | L | 6076 | 9.1 |
| SCon5 | Surgical Con | R | 7092 | 10.2 |
| | Surgical Con | L | 8644 | 9.8 |
| SCon6 | Surgical Con | R | 7035 | 10.8 |
| | Surgical Con | L | 8574 | 9.9 |
| PDX1 | Tumor | R | 5817 | 7 |
| PDX2 | Tumor | R | 7271 | 8.8 |
| PDX3 | Tumor | R | 4975 | 8.8 |
| PDX4 | Tumor | R | 6886 | 8.5 |
| PDX5 | Tumor | R | 6649 | 6.3 |
| PDX6 | Tumor | R | 7264 | 8 |
