## Supplemental Table 2 for "Pioglitazone administration restores a PPARG-dependent transcriptional network and ATP levels within skeletal muscles of mice implanted with patient-derived breast tumors"

**Supplemental Table 2: Differentially expressed genes in muscles of BC-PDOX mice compared to NSG-Con.**

| GeneID | Gene | baseMean | stat | log2FoldChange | pvalue | padj |
|---|---|---|---|---|---|---|
| ENSMUSG00000000088 | Cox5a | 3167.84 | -4.1921 | -0.4446 | 0.0000 | 0.0033 |
| ENSMUSG00000000305 | Cdh4 | 371.62 | -3.9166 | -1.1030 | 0.0001 | 0.0070 |
| ENSMUSG00000000318 | Clec10a | 83.87 | 4.2462 | 1.9226 | 0.0000 | 0.0028 |
| ENSMUSG00000000399 | Ndufa9 | 2534.47 | -3.5744 | -0.3529 | 0.0004 | 0.0168 |
| ENSMUSG00000000794 | Kcnn3 | 103.10 | 3.2945 | 0.7882 | 0.0010 | 0.0328 |
| ENSMUSG00000000934 | Top1mt | 149.80 | -3.1864 | -0.5362 | 0.0014 | 0.0411 |
| ENSMUSG00000001175 | Calm1 | 10367.67 | -5.1854 | -0.4562 | 0.0000 | 0.0001 |
| ENSMUSG00000001227 | Sema6b | 80.51 | 4.3431 | 1.0456 | 0.0000 | 0.0020 |
| ENSMUSG00000001604 | Tcea3 | 1802.63 | -3.8036 | -0.3941 | 0.0001 | 0.0094 |
| ENSMUSG00000001930 | Vwf | 682.89 | 3.8075 | 0.6450 | 0.0001 | 0.0093 |
| ENSMUSG00000002010 | Idh3g | 2826.85 | -4.4065 | -0.3741 | 0.0000 | 0.0017 |
| ENSMUSG00000002012 | Pnck | 58.77 | -3.2019 | -0.9277 | 0.0014 | 0.0400 |
| ENSMUSG00000002379 | Ndufa11 | 936.15 | -3.2809 | -0.3485 | 0.0010 | 0.0339 |
| ENSMUSG00000002416 | Ndufb2 | 908.37 | -3.9479 | -0.5043 | 0.0001 | 0.0065 |
| ENSMUSG00000002699 | Lcp2 | 19.13 | 3.3702 | 1.9270 | 0.0008 | 0.0279 |
| ENSMUSG00000003226 | Ranbp2 | 1668.60 | 3.5802 | 0.3594 | 0.0003 | 0.0165 |
| ENSMUSG00000003283 | Hck | 9.18 | 3.1762 | 2.1417 | 0.0015 | 0.0419 |
| ENSMUSG00000003541 | Ier3 | 85.16 | 3.5340 | 0.7586 | 0.0004 | 0.0185 |
| ENSMUSG00000003746 | Man1a | 372.27 | 3.2103 | 0.6709 | 0.0013 | 0.0391 |
| ENSMUSG00000003848 | Nob1 | 217.97 | -3.3298 | -0.4765 | 0.0009 | 0.0303 |
| ENSMUSG00000004040 | Stat3 | 1781.13 | 12.480 | 1.3954 | 0.0000 | 0.0000 |
| ENSMUSG00000004319 | Clcn3 | 1011.91 | 3.1180 | 0.2672 | 0.0018 | 0.0471 |
| ENSMUSG00000004609 | Cd33 | 54.35 | 3.3181 | 1.0379 | 0.0009 | 0.0310 |
| ENSMUSG00000004610 | Etfb | 1587.08 | -3.7500 | -0.3661 | 0.0002 | 0.0109 |
| ENSMUSG00000004730 | Adgre1 | 54.61 | 3.3381 | 1.6106 | 0.0008 | 0.0298 |
| ENSMUSG00000004891 | Nes | 459.52 | -3.7886 | -0.5211 | 0.0002 | 0.0097 |
| ENSMUSG00000004939 | Nmrk2 | 511.01 | -5.3098 | -2.3492 | 0.0000 | 0.0000 |
| ENSMUSG00000005268 | Prlr | 31.65 | 3.3717 | 1.9251 | 0.0007 | 0.0278 |
| ENSMUSG00000005299 | Letm1 | 788.85 | -3.1185 | -0.2866 | 0.0018 | 0.0471 |
| ENSMUSG00000005373 | Mlxipl | 1222.57 | -3.1571 | -0.4593 | 0.0016 | 0.0434 |
| ENSMUSG00000005378 | Bud23 | 155.15 | -3.2515 | -0.4673 | 0.0011 | 0.0360 |
| ENSMUSG00000005510 | Ndufs3 | 2956.16 | -4.9628 | -0.4529 | 0.0000 | 0.0002 |
| ENSMUSG00000005580 | Adcy9 | 627.19 | 3.6447 | 0.4552 | 0.0003 | 0.0140 |
| ENSMUSG00000005871 | Apc | 1090.75 | 3.3112 | 0.3620 | 0.0009 | 0.0315 |
| ENSMUSG00000005981 | Trap1 | 1499.98 | -3.3474 | -0.2765 | 0.0008 | 0.0292 |
| ENSMUSG00000006057 | Atp5g1 | 2916.29 | -3.6763 | -0.3857 | 0.0002 | 0.0129 |
| ENSMUSG00000006281 | Tep1 | 371.22 | 3.3157 | 0.3395 | 0.0009 | 0.0311 |
| ENSMUSG00000006344 | Ggt5 | 97.51 | 3.2581 | 0.9108 | 0.0011 | 0.0356 |
| ENSMUSG00000008348 | Ubc | 13376.21 | -3.1514 | -0.6960 | 0.0016 | 0.0439 |
| ENSMUSG00000008496 | Pou2f2 | 24.60 | 3.7231 | 1.8021 | 0.0002 | 0.0117 |
| ENSMUSG00000008540 | Mgst1 | 255.74 | 3.2597 | 0.8797 | 0.0011 | 0.0355 |
| ENSMUSG00000009185 | Ccl8 | 134.01 | 4.7870 | 4.9934 | 0.0000 | 0.0004 |
| ENSMUSG00000009376 | Met | 321.39 | 3.7063 | 0.6382 | 0.0002 | 0.0123 |
| ENSMUSG00000009647 | Mcu | 873.30 | 7.2021 | 0.6023 | 0.0000 | 0.0000 |

| | | | | | | |
|---|---|---|---|---|---|---|
| ENSMUSG00000009863 | Sdhb | 3900.47 | -4.1852 | -0.4860 | 0.0000 | 0.0033 |
| ENSMUSG00000011263 | Exoc3l2 | 22.92 | 3.0950 | 1.3905 | 0.0020 | 0.0495 |
| ENSMUSG00000013584 | Aldh1a2 | 100.55 | 3.8806 | 1.4612 | 0.0001 | 0.0079 |
| ENSMUSG00000014294 | Ndufa2 | 872.91 | -3.1925 | -0.3775 | 0.0014 | 0.0407 |
| ENSMUSG00000014313 | Cox6c | 3252.61 | -3.2611 | -0.3283 | 0.0011 | 0.0354 |
| ENSMUSG00000014355 | Anapc1 | 803.03 | 3.7770 | 0.3493 | 0.0002 | 0.0100 |
| ENSMUSG00000014444 | Piezo1 | 893.37 | 6.2560 | 1.2649 | 0.0000 | 0.0000 |
| ENSMUSG00000015087 | Rabl6 | 1361.81 | -3.6913 | -0.3722 | 0.0002 | 0.0127 |
| ENSMUSG00000015143 | Actn1 | 255.73 | 4.1996 | 0.5976 | 0.0000 | 0.0033 |
| ENSMUSG00000015243 | Abca1 | 721.65 | 4.3453 | 0.9528 | 0.0000 | 0.0020 |
| ENSMUSG00000015340 | Cybb | 71.35 | 3.4818 | 1.1536 | 0.0005 | 0.0212 |
| ENSMUSG00000015451 | C4a | 53.57 | 3.1098 | 1.2678 | 0.0019 | 0.0479 |
| ENSMUSG00000015745 | Plekho1 | 119.42 | 3.2247 | 0.5725 | 0.0013 | 0.0376 |
| ENSMUSG00000015766 | Eps8 | 113.90 | 3.1054 | 0.8521 | 0.0019 | 0.0484 |
| ENSMUSG00000015843 | Rxrg | 816.06 | -3.7376 | -0.4770 | 0.0002 | 0.0112 |
| ENSMUSG00000015850 | Adamtsl4 | 423.30 | 3.5939 | 0.6582 | 0.0003 | 0.0160 |
| ENSMUSG00000015852 | Fcrls | 42.27 | 3.3736 | 1.4548 | 0.0007 | 0.0277 |
| ENSMUSG00000016252 | Atp5e | 2992.87 | -3.4693 | -0.3843 | 0.0005 | 0.0219 |
| ENSMUSG00000016427 | Ndufa1 | 785.72 | -3.8022 | -0.3724 | 0.0001 | 0.0094 |
| ENSMUSG00000016534 | Lamp2 | 1778.48 | 4.4476 | 0.4683 | 0.0000 | 0.0015 |
| ENSMUSG00000016756 | Cmah | 120.23 | 6.6363 | 1.5105 | 0.0000 | 0.0000 |
| ENSMUSG00000016921 | Srsf6 | 940.00 | -3.3278 | -0.3080 | 0.0009 | 0.0303 |
| ENSMUSG00000017300 | Tnnc2 | 82740.97 | -3.3894 | -0.4711 | 0.0007 | 0.0267 |
| ENSMUSG00000017677 | Wsb1 | 187.64 | -3.3358 | -0.5497 | 0.0009 | 0.0298 |
| ENSMUSG00000017774 | Myo1c | 887.20 | 3.2320 | 0.3354 | 0.0012 | 0.0370 |
| ENSMUSG00000017778 | Cox7c | 3169.56 | -3.2965 | -0.3522 | 0.0010 | 0.0327 |
| ENSMUSG00000017781 | Pitpna | 2020.79 | -3.5211 | -0.6691 | 0.0004 | 0.0190 |
| ENSMUSG00000018008 | Cyth4 | 77.73 | 3.7427 | 1.5686 | 0.0002 | 0.0111 |
| ENSMUSG00000018340 | Anxa6 | 2504.20 | 4.7420 | 0.5752 | 0.0000 | 0.0005 |
| ENSMUSG00000018446 | C1qbp | 681.91 | -3.1957 | -0.3046 | 0.0014 | 0.0404 |
| ENSMUSG00000018554 | Ybx2 | 101.97 | -4.2524 | -0.8422 | 0.0000 | 0.0028 |
| ENSMUSG00000018654 | Ikzf1 | 18.44 | 3.4566 | 1.9375 | 0.0005 | 0.0228 |
| ENSMUSG00000018770 | Atp5g3 | 4938.67 | -4.0281 | -0.4166 | 0.0001 | 0.0051 |
| ENSMUSG00000018800 | Abca5 | 490.55 | 3.9847 | 0.6713 | 0.0001 | 0.0057 |
| ENSMUSG00000018846 | Pank3 | 612.28 | 3.8052 | 0.5852 | 0.0001 | 0.0093 |
| ENSMUSG00000018927 | Ccl6 | 153.25 | 3.6651 | 2.3312 | 0.0002 | 0.0132 |
| ENSMUSG00000019122 | Ccl9 | 96.78 | 4.7181 | 1.9731 | 0.0000 | 0.0006 |
| ENSMUSG00000019179 | Mdh2 | 7280.34 | -4.1556 | -0.4256 | 0.0000 | 0.0035 |
| ENSMUSG00000019590 | Cyb561 | 77.03 | 3.7284 | 1.0289 | 0.0002 | 0.0115 |
| ENSMUSG00000020017 | Hal | 122.93 | 5.1323 | 1.8436 | 0.0000 | 0.0001 |
| ENSMUSG00000020022 | Ndufa12 | 1720.46 | -3.3675 | -0.3129 | 0.0008 | 0.0280 |
| ENSMUSG00000020027 | Socs2 | 532.90 | -4.7636 | -1.3152 | 0.0000 | 0.0005 |
| ENSMUSG00000020057 | Dram1 | 73.99 | 3.4019 | 1.2156 | 0.0007 | 0.0261 |
| ENSMUSG00000020099 | Unc5b | 58.99 | -3.3157 | -0.8686 | 0.0009 | 0.0311 |
| ENSMUSG00000020108 | Ddit4 | 109.28 | 4.3327 | 1.4448 | 0.0000 | 0.0021 |
| ENSMUSG00000020153 | Ndufs7 | 1850.07 | -4.5145 | -0.4825 | 0.0000 | 0.0012 |
| ENSMUSG00000020163 | Uqcr11 | 1983.89 | -3.9541 | -0.4637 | 0.0001 | 0.0064 |
| ENSMUSG00000020227 | Irak3 | 86.30 | 4.2317 | 1.0283 | 0.0000 | 0.0029 |

| | | | | | | |
|---|---|---|---|---|---|---|
| ENSMUSG00000020354 | Sgcd | 520.07 | 4.9996 | 1.0831 | 0.0000 | 0.0002 |
| ENSMUSG00000020475 | Pgam2 | 15669.98 | -3.4955 | -0.5180 | 0.0005 | 0.0203 |
| ENSMUSG00000020483 | Dynll2 | 3290.64 | -3.6771 | -0.5795 | 0.0002 | 0.0129 |
| ENSMUSG00000020532 | Acaca | 663.27 | 4.0546 | 0.8727 | 0.0001 | 0.0046 |
| ENSMUSG00000020681 | Ace | 677.21 | 5.0177 | 0.9958 | 0.0000 | 0.0002 |
| ENSMUSG00000020717 | Pecam1 | 1006.68 | 3.6742 | 0.4028 | 0.0002 | 0.0129 |
| ENSMUSG00000020744 | Slc25a19 | 367.96 | -3.2008 | -0.3677 | 0.0014 | 0.0400 |
| ENSMUSG00000020841 | Cpd | 1683.83 | 3.1956 | 0.3793 | 0.0014 | 0.0404 |
| ENSMUSG00000020846 | Rflnb | 277.30 | -4.2669 | -0.7636 | 0.0000 | 0.0027 |
| ENSMUSG00000021036 | Sptlc2 | 149.20 | 3.2978 | 0.5483 | 0.0010 | 0.0326 |
| ENSMUSG00000021091 | Serpina3n | 286.00 | 4.1935 | 3.2185 | 0.0000 | 0.0033 |
| ENSMUSG00000021109 | Hif1a | 482.08 | 3.2200 | 0.4768 | 0.0013 | 0.0380 |
| ENSMUSG00000021190 | Lgmn | 473.22 | 3.2926 | 0.8130 | 0.0010 | 0.0328 |
| ENSMUSG00000021200 | Asb2 | 5604.25 | -3.1015 | -0.5305 | 0.0019 | 0.0488 |
| ENSMUSG00000021235 | Coq6 | 167.93 | -3.1933 | -0.4411 | 0.0014 | 0.0406 |
| ENSMUSG00000021280 | Exoc3l4 | 75.49 | 4.1276 | 1.9831 | 0.0000 | 0.0037 |
| ENSMUSG00000021281 | Tnfaip2 | 1565.88 | 4.5177 | 0.9581 | 0.0000 | 0.0012 |
| ENSMUSG00000021290 | Atp5mpl | 1516.69 | -4.6303 | -0.4518 | 0.0000 | 0.0008 |
| ENSMUSG00000021423 | Ly86 | 19.44 | 4.6991 | 2.7887 | 0.0000 | 0.0006 |
| ENSMUSG00000021520 | Uqcrb | 3274.70 | -3.4531 | -0.3384 | 0.0006 | 0.0230 |
| ENSMUSG00000021596 | Mctp1 | 40.47 | 3.5343 | 1.1157 | 0.0004 | 0.0185 |
| ENSMUSG00000021606 | Ndufs6 | 931.95 | -3.2962 | -0.3116 | 0.0010 | 0.0327 |
| ENSMUSG00000021710 | Nln | 117.33 | 3.3649 | 0.6926 | 0.0008 | 0.0281 |
| ENSMUSG00000021771 | Vdac2 | 3351.45 | -3.1963 | -0.2261 | 0.0014 | 0.0404 |
| ENSMUSG00000021796 | Bmpr1a | 1561.07 | 4.0743 | 0.3383 | 0.0000 | 0.0043 |
| ENSMUSG00000021814 | Anxa7 | 1006.49 | 3.1687 | 0.4911 | 0.0015 | 0.0426 |
| ENSMUSG00000021898 | Asb14 | 1184.36 | -3.1344 | -0.2628 | 0.0017 | 0.0455 |
| ENSMUSG00000021903 | Galnt15 | 652.80 | 4.5826 | 1.0733 | 0.0000 | 0.0009 |
| ENSMUSG00000021950 | Anxa8 | 27.11 | 3.6242 | 2.3887 | 0.0003 | 0.0148 |
| ENSMUSG00000021983 | Atp8a2\|LOC108168164 | 116.24 | 6.3293 | 1.3258 | 0.0000 | 0.0000 |
| ENSMUSG00000021993 | Mipep | 492.75 | -3.2427 | -0.3773 | 0.0012 | 0.0369 |
| ENSMUSG00000021998 | Lcp1 | 108.17 | 3.5330 | 1.4115 | 0.0004 | 0.0185 |
| ENSMUSG00000022013 | Dnajc15 | 502.56 | -3.5498 | -0.3924 | 0.0004 | 0.0179 |
| ENSMUSG00000022037 | Clu | 455.59 | 3.3838 | 0.6347 | 0.0007 | 0.0271 |
| ENSMUSG00000022146 | Osmr | 263.68 | 6.6125 | 1.2626 | 0.0000 | 0.0000 |
| ENSMUSG00000022148 | Fyb | 32.18 | 3.4396 | 1.4506 | 0.0006 | 0.0238 |
| ENSMUSG00000022150 | Dab2 | 429.91 | 3.5816 | 0.9541 | 0.0003 | 0.0165 |
| ENSMUSG00000022181 | C6 | 27.35 | 4.9326 | 4.7721 | 0.0000 | 0.0002 |
| ENSMUSG00000022186 | Oxct1 | 3138.30 | -3.7781 | -0.5996 | 0.0002 | 0.0100 |
| ENSMUSG00000022354 | Ndufb9 | 3182.13 | -4.2016 | -0.4155 | 0.0000 | 0.0033 |
| ENSMUSG00000022488 | Nckap1l | 47.14 | 3.4674 | 1.5131 | 0.0005 | 0.0220 |
| ENSMUSG00000022490 | Ppp1r1a | 463.13 | -3.1160 | -0.4036 | 0.0018 | 0.0472 |
| ENSMUSG00000022500 | Litaf\|Gm9861 | 128.16 | 3.6909 | 0.9043 | 0.0002 | 0.0127 |
| ENSMUSG00000022533 | Atp13a3 | 731.47 | 3.1754 | 0.3785 | 0.0015 | 0.0419 |
| ENSMUSG00000022551 | Cyc1 | 3307.54 | -3.5280 | -0.4299 | 0.0004 | 0.0187 |
| ENSMUSG00000022748 | Cmss1 | 210.11 | -3.3788 | -0.5178 | 0.0007 | 0.0274 |
| ENSMUSG00000022831 | Hcls1 | 37.72 | 3.3625 | 1.1951 | 0.0008 | 0.0282 |
| ENSMUSG00000022890 | Atp5j | 3687.51 | -4.2980 | -0.4574 | 0.0000 | 0.0024 |

| | | | | | | |
|---|---|---|---|---|---|---|
| ENSMUSG00000022895 | Ets2 | 937.01 | 5.6462 | 0.5401 | 0.0000 | 0.0000 |
| ENSMUSG00000022912 | Pros1 | 232.43 | 3.2073 | 0.7606 | 0.0013 | 0.0393 |
| ENSMUSG00000022952 | Runx1 | 91.14 | 4.8825 | 1.1992 | 0.0000 | 0.0003 |
| ENSMUSG00000022956 | Atp5o | 4072.59 | -4.0135 | -0.3823 | 0.0001 | 0.0053 |
| ENSMUSG00000023089 | Ndufa5 | 1382.43 | -4.4705 | -0.4277 | 0.0000 | 0.0014 |
| ENSMUSG00000023175 | Bsg | 6847.23 | -3.3209 | -0.3261 | 0.0009 | 0.0309 |
| ENSMUSG00000023186 | Vwa5a | 582.70 | 5.4278 | 0.7409 | 0.0000 | 0.0000 |
| ENSMUSG00000023367 | Tmem176a | 144.96 | 5.8453 | 1.6547 | 0.0000 | 0.0000 |
| ENSMUSG00000023830 | Igf2r | 2121.09 | 3.1691 | 0.2771 | 0.0015 | 0.0426 |
| ENSMUSG00000023913 | Pla2g7 | 673.62 | -3.4224 | -1.3333 | 0.0006 | 0.0248 |
| ENSMUSG00000023951 | Vegfa | 3075.70 | -3.2645 | -0.4956 | 0.0011 | 0.0351 |
| ENSMUSG00000024033 | Rsph1 | 57.71 | -5.7017 | -2.1114 | 0.0000 | 0.0000 |
| ENSMUSG00000024038 | Ndufv3 | 924.59 | -4.2309 | -0.4601 | 0.0000 | 0.0029 |
| ENSMUSG00000024059 | Clip4 | 1464.73 | 3.2378 | 0.7373 | 0.0012 | 0.0370 |
| ENSMUSG00000024082 | Ndufaf7 | 187.88 | -3.2691 | -0.5086 | 0.0011 | 0.0348 |
| ENSMUSG00000024085 | Man2a1 | 273.46 | 3.2877 | 0.6285 | 0.0010 | 0.0332 |
| ENSMUSG00000024099 | Ndufv2 | 3148.65 | -3.9869 | -0.4222 | 0.0001 | 0.0057 |
| ENSMUSG00000024168 | Tmem204 | 104.64 | -3.2689 | -0.8228 | 0.0011 | 0.0348 |
| ENSMUSG00000024180 | Tmem8 | 38.12 | 4.1465 | 2.1405 | 0.0000 | 0.0036 |
| ENSMUSG00000024194 | Cuta | 359.69 | -3.3816 | -0.4505 | 0.0007 | 0.0272 |
| ENSMUSG00000024247 | Pkdcc | 795.97 | -3.6769 | -0.5612 | 0.0002 | 0.0129 |
| ENSMUSG00000024294 | Mib1 | 1185.91 | -4.0853 | -0.5246 | 0.0000 | 0.0042 |
| ENSMUSG00000024440 | Pcdh12 | 89.40 | -3.7953 | -0.9261 | 0.0001 | 0.0095 |
| ENSMUSG00000024529 | Lox | 177.77 | 3.5833 | 1.1426 | 0.0003 | 0.0165 |
| ENSMUSG00000024576 | Csnk1a1 | 3719.65 | 3.3736 | 0.2429 | 0.0007 | 0.0277 |
| ENSMUSG00000024589 | Nedd4l | 813.94 | 7.8681 | 1.0056 | 0.0000 | 0.0000 |
| ENSMUSG00000024597 | Slc12a2 | 1961.34 | 5.0701 | 0.5903 | 0.0000 | 0.0001 |
| ENSMUSG00000024679 | Ms4a6d | 29.51 | 4.7160 | 2.1308 | 0.0000 | 0.0006 |
| ENSMUSG00000024683 | Mrpl16 | 395.24 | -3.5394 | -0.4045 | 0.0004 | 0.0182 |
| ENSMUSG00000024687 | Osbp | 1691.58 | 3.8582 | 0.3277 | 0.0001 | 0.0082 |
| ENSMUSG00000024962 | Vegfb | 972.81 | -3.4113 | -0.3735 | 0.0006 | 0.0255 |
| ENSMUSG00000024997 | Prdx3 | 1977.20 | -4.1696 | -0.3289 | 0.0000 | 0.0034 |
| ENSMUSG00000025017 | Pik3ap1 | 28.46 | 3.3084 | 1.6089 | 0.0009 | 0.0317 |
| ENSMUSG00000025035 | Arl3 | 501.59 | -3.3368 | -0.5466 | 0.0008 | 0.0298 |
| ENSMUSG00000025204 | Ndufb8 | 3070.78 | -3.6536 | -0.3804 | 0.0003 | 0.0137 |
| ENSMUSG00000025213 | Kazald1 | 70.56 | -3.1288 | -0.8520 | 0.0018 | 0.0460 |
| ENSMUSG00000025255 | Zfhx4 | 483.98 | 3.8965 | 0.3960 | 0.0001 | 0.0075 |
| ENSMUSG00000025261 | Huwe1 | 5475.64 | 3.3766 | 0.2715 | 0.0007 | 0.0276 |
| ENSMUSG00000025314 | Ptprj | 144.94 | 3.1192 | 0.8205 | 0.0018 | 0.0471 |
| ENSMUSG00000025348 | Itga7 | 1530.05 | 3.1440 | 0.5371 | 0.0017 | 0.0447 |
| ENSMUSG00000025488 | Cox8b | 2310.10 | -4.3459 | -0.5303 | 0.0000 | 0.0020 |
| ENSMUSG00000025492 | Ifitm3 | 381.95 | 4.1225 | 0.6762 | 0.0000 | 0.0038 |
| ENSMUSG00000025511 | Tspan4 | 102.67 | 3.4441 | 0.7882 | 0.0006 | 0.0236 |
| ENSMUSG00000025586 | Cpeb1 | 628.77 | -3.3529 | -0.7378 | 0.0008 | 0.0287 |
| ENSMUSG00000025651 | Uqcrc1 | 5878.46 | -4.3664 | -0.4407 | 0.0000 | 0.0019 |
| ENSMUSG00000025701 | Alox5 | 19.97 | 3.6925 | 1.9065 | 0.0002 | 0.0127 |
| ENSMUSG00000025777 | Gdap1 | 262.69 | -4.2861 | -0.9404 | 0.0000 | 0.0025 |
| ENSMUSG00000025825 | Iscu | 872.44 | -3.4038 | -0.3876 | 0.0007 | 0.0260 |

| | | | | | | |
|---|---|---|---|---|---|---|
| ENSMUSG00000025887 | Casp12 | 174.86 | 3.5423 | 0.7655 | 0.0004 | 0.0182 |
| ENSMUSG00000025911 | Adhfe1 | 345.10 | -3.4010 | -0.3607 | 0.0007 | 0.0261 |
| ENSMUSG00000025950 | Idh1 | 365.12 | 3.3045 | 0.6640 | 0.0010 | 0.0321 |
| ENSMUSG00000026032 | Ndufb3 | 992.76 | -3.4362 | -0.3548 | 0.0006 | 0.0240 |
| ENSMUSG00000026062 | Slc9a2 | 501.61 | -3.1008 | -0.3533 | 0.0019 | 0.0489 |
| ENSMUSG00000026072 | Il1r1 | 166.39 | 3.1991 | 0.7064 | 0.0014 | 0.0402 |
| ENSMUSG00000026260 | Ndufa10 | 2856.22 | -4.2140 | -0.3915 | 0.0000 | 0.0031 |
| ENSMUSG00000026321 | Tnfrsf11a | 33.06 | 3.1127 | 1.2917 | 0.0019 | 0.0475 |
| ENSMUSG00000026395 | Ptprc | 52.67 | 3.3270 | 1.2959 | 0.0009 | 0.0303 |
| ENSMUSG00000026436 | Elk4 | 936.42 | 3.5560 | 0.3464 | 0.0004 | 0.0177 |
| ENSMUSG00000026480 | Ncf2 | 26.81 | 3.2682 | 1.3341 | 0.0011 | 0.0348 |
| ENSMUSG00000026526 | Fh1 | 2301.37 | -4.3709 | -0.3819 | 0.0000 | 0.0019 |
| ENSMUSG00000026568 | Mpc2 | 1919.13 | -4.1043 | -0.4186 | 0.0000 | 0.0040 |
| ENSMUSG00000026580 | Selp | 81.23 | 4.7503 | 1.3690 | 0.0000 | 0.0005 |
| ENSMUSG00000026589 | Sec16b | 48.71 | 3.2631 | 0.9330 | 0.0011 | 0.0352 |
| ENSMUSG00000026656 | Fcgr2b | 58.77 | 3.2558 | 1.3768 | 0.0011 | 0.0358 |
| ENSMUSG00000026712 | Mrc1 | 348.88 | 3.6429 | 1.7530 | 0.0003 | 0.0140 |
| ENSMUSG00000026822 | Lcn2 | 53.00 | 3.3381 | 3.6911 | 0.0008 | 0.0298 |
| ENSMUSG00000026895 | Ndufa8 | 1679.16 | -3.3504 | -0.3536 | 0.0008 | 0.0289 |
| ENSMUSG00000027322 | Siglec1 | 63.41 | 4.4633 | 2.2697 | 0.0000 | 0.0014 |
| ENSMUSG00000027332 | Ivd | 1137.73 | -3.4261 | -0.4567 | 0.0006 | 0.0245 |
| ENSMUSG00000027371 | Fahd2a | 170.07 | -4.1965 | -0.6174 | 0.0000 | 0.0033 |
| ENSMUSG00000027406 | Idh3b | 4573.34 | -4.1597 | -0.3823 | 0.0000 | 0.0034 |
| ENSMUSG00000027546 | Atp9a | 954.53 | 7.6373 | 1.5421 | 0.0000 | 0.0000 |
| ENSMUSG00000027624 | Epb41l1 | 151.72 | 3.2114 | 0.6694 | 0.0013 | 0.0390 |
| ENSMUSG00000027634 | Ndrg3 | 631.48 | 4.3171 | 0.4000 | 0.0000 | 0.0022 |
| ENSMUSG00000028028 | Alpk1 | 132.06 | 3.1303 | 0.9586 | 0.0017 | 0.0460 |
| ENSMUSG00000028195 | Cyr61 | 485.31 | 5.8016 | 0.8878 | 0.0000 | 0.0000 |
| ENSMUSG00000028197 | Col24a1 | 93.53 | -6.0829 | -1.4848 | 0.0000 | 0.0000 |
| ENSMUSG00000028211 | Trp53inp1 | 285.38 | 3.1399 | 0.4798 | 0.0017 | 0.0451 |
| ENSMUSG00000028381 | Ugcg | 235.68 | 3.2533 | 0.5883 | 0.0011 | 0.0360 |
| ENSMUSG00000028405 | Aco1 | 300.69 | 3.7012 | 0.4954 | 0.0002 | 0.0124 |
| ENSMUSG00000028420 | Tmem38b | 1580.80 | 3.3276 | 0.7944 | 0.0009 | 0.0303 |
| ENSMUSG00000028444 | Cntfr | 203.81 | -4.1098 | -0.7945 | 0.0000 | 0.0039 |
| ENSMUSG00000028518 | Prkaa2 | 7428.62 | 3.5895 | 0.4378 | 0.0003 | 0.0161 |
| ENSMUSG00000028581 | Laptm5 | 141.20 | 3.5528 | 1.2987 | 0.0004 | 0.0178 |
| ENSMUSG00000028600 | Podn | 181.20 | 3.5665 | 0.8697 | 0.0004 | 0.0172 |
| ENSMUSG00000028630 | Dyrk2 | 1970.17 | -3.9134 | -0.5848 | 0.0001 | 0.0071 |
| ENSMUSG00000028648 | Ndufs5 | 809.66 | -4.1618 | -0.4011 | 0.0000 | 0.0034 |
| ENSMUSG00000028737 | Aldh4a1 | 1196.22 | -3.1752 | -0.3468 | 0.0015 | 0.0419 |
| ENSMUSG00000028862 | Map3k6 | 59.91 | 3.7395 | 1.2995 | 0.0002 | 0.0112 |
| ENSMUSG00000028944 | Prkag2 | 110.77 | 3.1605 | 0.6617 | 0.0016 | 0.0431 |
| ENSMUSG00000028949 | Smarcd3 | 1637.80 | -4.8119 | -0.4799 | 0.0000 | 0.0004 |
| ENSMUSG00000029082 | Bst1 | 11.65 | 3.6624 | 2.3750 | 0.0002 | 0.0133 |
| ENSMUSG00000029287 | Tgfbr3 | 538.04 | 3.6840 | 0.7196 | 0.0002 | 0.0129 |
| ENSMUSG00000029321 | Slc10a6 | 16.87 | 3.6896 | 2.3294 | 0.0002 | 0.0127 |
| ENSMUSG00000029471 | Camkk2 | 104.64 | 3.5737 | 0.7016 | 0.0004 | 0.0168 |
| ENSMUSG00000029559 | 2210016L21Rik | 158.88 | -3.1074 | -0.5306 | 0.0019 | 0.0482 |

| | | | | | | |
|---|---|---|---|---|---|---|
| ENSMUSG00000029632 | Ndufa4 | 3594.66 | -3.4047 | -0.3651 | 0.0007 | 0.0260 |
| ENSMUSG00000029810 | Tmem176b | 218.99 | 4.8243 | 1.2250 | 0.0000 | 0.0004 |
| ENSMUSG00000029864 | Gstk1 | 261.51 | -3.1622 | -0.5515 | 0.0016 | 0.0430 |
| ENSMUSG00000029994 | Anxa4 | 315.07 | 3.6676 | 0.7531 | 0.0002 | 0.0131 |
| ENSMUSG00000030043 | Tacr1 | 18.34 | 4.5070 | 3.1739 | 0.0000 | 0.0012 |
| ENSMUSG00000030089 | Slc41a3 | 1047.79 | -3.4114 | -0.6583 | 0.0006 | 0.0255 |
| ENSMUSG00000030095 | Tmem43 | 405.34 | 3.5922 | 0.4601 | 0.0003 | 0.0161 |
| ENSMUSG00000030213 | Atf7ip | 1039.74 | 4.7476 | 0.4552 | 0.0000 | 0.0005 |
| ENSMUSG00000030341 | Tnfrsf1a | 311.44 | 4.2774 | 0.8621 | 0.0000 | 0.0026 |
| ENSMUSG00000030579 | Tyrobp | 34.82 | 3.2928 | 1.4228 | 0.0010 | 0.0328 |
| ENSMUSG00000030737 | Slco2b1 | 208.30 | 4.4004 | 0.9211 | 0.0000 | 0.0017 |
| ENSMUSG00000030748 | Il4ra | 227.31 | 3.5456 | 1.1738 | 0.0004 | 0.0181 |
| ENSMUSG00000030772 | Dkk3 | 76.49 | -3.1183 | -0.9976 | 0.0018 | 0.0471 |
| ENSMUSG00000030785 | Cox6a2 | 8457.47 | -3.8442 | -0.4071 | 0.0001 | 0.0085 |
| ENSMUSG00000030786 | Itgam | 59.27 | 3.6163 | 1.2923 | 0.0003 | 0.0150 |
| ENSMUSG00000030793 | Pycard | 17.85 | 3.2337 | 1.5915 | 0.0012 | 0.0370 |
| ENSMUSG00000030798 | Cd37 | 24.69 | 3.6766 | 1.7968 | 0.0002 | 0.0129 |
| ENSMUSG00000030826 | Bcat2 | 553.11 | -4.4152 | -0.4431 | 0.0000 | 0.0016 |
| ENSMUSG00000030869 | Ndufab1 | 1497.71 | -3.4654 | -0.3385 | 0.0005 | 0.0221 |
| ENSMUSG00000030884 | Uqcrc2 | 4785.62 | -3.5306 | -0.3549 | 0.0004 | 0.0186 |
| ENSMUSG00000031010 | Usp9x | 4328.18 | 3.7632 | 0.2919 | 0.0002 | 0.0104 |
| ENSMUSG00000031027 | Stk33 | 275.44 | -7.2191 | -1.6195 | 0.0000 | 0.0000 |
| ENSMUSG00000031059 | Ndufb11 | 2127.91 | -3.3577 | -0.3570 | 0.0008 | 0.0284 |
| ENSMUSG00000031097 | Tnni2 | 61290.15 | -4.3109 | -0.3652 | 0.0000 | 0.0023 |
| ENSMUSG00000031129 | Slc9a9 | 52.56 | 3.6498 | 1.4100 | 0.0003 | 0.0138 |
| ENSMUSG00000031231 | Cox7b | 3512.64 | -3.7045 | -0.3762 | 0.0002 | 0.0123 |
| ENSMUSG00000031309 | Rps6ka3 | 1766.90 | 4.1850 | 0.4439 | 0.0000 | 0.0033 |
| ENSMUSG00000031373 | Car5b | 81.56 | 3.6681 | 1.5623 | 0.0002 | 0.0131 |
| ENSMUSG00000031453 | Rasa3 | 197.99 | 4.3517 | 0.7249 | 0.0000 | 0.0020 |
| ENSMUSG00000031494 | Cd209a | 26.05 | 3.2320 | 2.0031 | 0.0012 | 0.0370 |
| ENSMUSG00000031543 | Ank1 | 7549.35 | -3.8520 | -0.5278 | 0.0001 | 0.0084 |
| ENSMUSG00000031608 | Galnt7 | 101.49 | 3.8790 | 0.8437 | 0.0001 | 0.0079 |
| ENSMUSG00000031633 | Slc25a4 | 34248.29 | -3.8015 | -0.4284 | 0.0001 | 0.0094 |
| ENSMUSG00000031683 | Lsm6 | 483.64 | -3.1309 | -0.3197 | 0.0017 | 0.0459 |
| ENSMUSG00000031722 | Hp | 378.50 | 4.2620 | 4.1402 | 0.0000 | 0.0027 |
| ENSMUSG00000031762 | Mt2 | 216.16 | 4.0039 | 3.8654 | 0.0001 | 0.0054 |
| ENSMUSG00000031765 | Mt1 | 189.43 | 3.5498 | 3.1518 | 0.0004 | 0.0179 |
| ENSMUSG00000031803 | B3gnt3 | 88.17 | 5.3427 | 1.1229 | 0.0000 | 0.0000 |
| ENSMUSG00000031805 | Jak3 | 171.56 | 6.0830 | 0.9531 | 0.0000 | 0.0000 |
| ENSMUSG00000031818 | Cox4i1 | 5074.26 | -3.7857 | -0.3673 | 0.0002 | 0.0098 |
| ENSMUSG00000031958 | Ldhd | 413.87 | -3.1411 | -0.4699 | 0.0017 | 0.0450 |
| ENSMUSG00000031963 | Bmper | 100.42 | 3.2369 | 0.8430 | 0.0012 | 0.0370 |
| ENSMUSG00000031967 | Afg3l1 | 754.82 | -3.2319 | -0.3324 | 0.0012 | 0.0370 |
| ENSMUSG00000032013 | Trim29 | 9.16 | 4.0136 | 5.9041 | 0.0001 | 0.0053 |
| ENSMUSG00000032135 | Mcam | 342.03 | 3.1894 | 0.4659 | 0.0014 | 0.0408 |
| ENSMUSG00000032216 | Nedd4 | 10383.62 | 4.9843 | 0.3061 | 0.0000 | 0.0002 |
| ENSMUSG00000032239 | Rp9 | 259.19 | -3.6812 | -0.4632 | 0.0002 | 0.0129 |
| ENSMUSG00000032265 | Tent5a | 130.79 | 3.4536 | 0.9439 | 0.0006 | 0.0230 |

| | | | | | | |
|---|---|---|---|---|---|---|
| ENSMUSG00000032279 | Idh3a | 7077.37 | -3.2713 | -0.2673 | 0.0011 | 0.0347 |
| ENSMUSG00000032330 | Cox7a2 | 1239.02 | -3.1671 | -0.3414 | 0.0015 | 0.0426 |
| ENSMUSG00000032340 | Neo1 | 1095.32 | 5.3042 | 0.4409 | 0.0000 | 0.0000 |
| ENSMUSG00000032355 | Mlip | 1570.84 | 4.6625 | 0.3950 | 0.0000 | 0.0007 |
| ENSMUSG00000032403 | 2300009A05Rik | 82.30 | -3.1055 | -0.6204 | 0.0019 | 0.0484 |
| ENSMUSG00000032409 | Atr | 161.42 | 3.8262 | 0.5814 | 0.0001 | 0.0088 |
| ENSMUSG00000032420 | Nt5e | 102.73 | 3.1373 | 0.6483 | 0.0017 | 0.0454 |
| ENSMUSG00000032437 | Stt3b | 1983.61 | 3.5013 | 0.6650 | 0.0005 | 0.0200 |
| ENSMUSG00000032503 | Arpp21 | 200.60 | 3.1267 | 0.7372 | 0.0018 | 0.0461 |
| ENSMUSG00000032527 | Pccb | 875.10 | -3.5018 | -0.3646 | 0.0005 | 0.0200 |
| ENSMUSG00000032625 | Thsd7a | 84.99 | 3.1509 | 0.7183 | 0.0016 | 0.0439 |
| ENSMUSG00000032766 | Gng11 | 144.87 | -3.7912 | -0.6143 | 0.0001 | 0.0097 |
| ENSMUSG00000032892 | Rangrf | 301.14 | -3.1360 | -0.4009 | 0.0017 | 0.0454 |
| ENSMUSG00000032897 | Nfyc | 409.87 | -3.4773 | -0.3845 | 0.0005 | 0.0215 |
| ENSMUSG00000033044 | Dhrs7c | 2113.90 | -3.1448 | -0.3730 | 0.0017 | 0.0447 |
| ENSMUSG00000033161 | Atp1a1 | 1184.37 | -3.9417 | -0.5215 | 0.0001 | 0.0065 |
| ENSMUSG00000033671 | Cep350 | 1009.11 | 3.3915 | 0.3794 | 0.0007 | 0.0267 |
| ENSMUSG00000033685 | Ucp2 | 161.56 | 3.7289 | 1.1372 | 0.0002 | 0.0115 |
| ENSMUSG00000033788 | Dysf | 2466.38 | 3.7146 | 0.4019 | 0.0002 | 0.0120 |
| ENSMUSG00000033792 | Atp7a | 130.16 | 3.8265 | 0.7261 | 0.0001 | 0.0088 |
| ENSMUSG00000033938 | Ndufb7 | 1234.39 | -3.6171 | -0.4910 | 0.0003 | 0.0150 |
| ENSMUSG00000033965 | Slc16a2 | 154.15 | 3.8731 | 0.9135 | 0.0001 | 0.0079 |
| ENSMUSG00000034021 | Pds5b | 1039.82 | 3.8279 | 0.3500 | 0.0001 | 0.0088 |
| ENSMUSG00000034040 | Galnt17 | 173.35 | 6.7680 | 1.5141 | 0.0000 | 0.0000 |
| ENSMUSG00000034116 | Vav1 | 16.03 | 3.6816 | 2.3196 | 0.0002 | 0.0129 |
| ENSMUSG00000034254 | Agpat1 | 1073.45 | -4.2297 | -0.3139 | 0.0000 | 0.0029 |
| ENSMUSG00000034422 | Parp14 | 209.38 | 3.5441 | 0.5334 | 0.0004 | 0.0182 |
| ENSMUSG00000034485 | Uaca | 3360.55 | -3.3104 | -0.3535 | 0.0009 | 0.0315 |
| ENSMUSG00000034566 | Atp5h | 2580.68 | -3.5958 | -0.3899 | 0.0003 | 0.0160 |
| ENSMUSG00000034593 | Myo5a | 257.43 | 3.7280 | 0.6783 | 0.0002 | 0.0115 |
| ENSMUSG00000034612 | Chst11 | 94.63 | 4.1844 | 1.5002 | 0.0000 | 0.0033 |
| ENSMUSG00000034616 | Ssh3 | 65.76 | 3.3649 | 0.7526 | 0.0008 | 0.0281 |
| ENSMUSG00000034648 | Lrrn1 | 107.31 | 3.4942 | 1.0412 | 0.0005 | 0.0203 |
| ENSMUSG00000034684 | Sema3f | 144.15 | 3.6531 | 0.6935 | 0.0003 | 0.0137 |
| ENSMUSG00000034845 | Plvap | 380.94 | 5.0135 | 0.9655 | 0.0000 | 0.0002 |
| ENSMUSG00000034932 | Mrpl54 | 283.98 | -3.7902 | -0.4754 | 0.0002 | 0.0097 |
| ENSMUSG00000034947 | Tmem106a | 47.95 | 3.2667 | 1.1959 | 0.0011 | 0.0349 |
| ENSMUSG00000035048 | Anapc13 | 409.79 | -3.3175 | -0.3996 | 0.0009 | 0.0310 |
| ENSMUSG00000035247 | Hectd1 | 3758.36 | 3.4281 | 0.3602 | 0.0006 | 0.0244 |
| ENSMUSG00000035284 | Vps13c | 810.22 | 3.8051 | 0.5275 | 0.0001 | 0.0093 |
| ENSMUSG00000035673 | Sbno2 | 249.74 | 3.6438 | 0.7850 | 0.0003 | 0.0140 |
| ENSMUSG00000035674 | Ndufa3 | 1067.26 | -3.6755 | -0.4053 | 0.0002 | 0.0129 |
| ENSMUSG00000035697 | Arhgap45 | 44.87 | 3.2994 | 1.2940 | 0.0010 | 0.0325 |
| ENSMUSG00000035778 | Ggta1 | 165.01 | 3.2782 | 0.5335 | 0.0010 | 0.0341 |
| ENSMUSG00000036199 | Ndufa13 | 2626.37 | -3.7097 | -0.4386 | 0.0002 | 0.0122 |
| ENSMUSG00000036241 | Ube2r2 | 2528.56 | -3.3463 | -0.4348 | 0.0008 | 0.0292 |
| ENSMUSG00000036256 | Igfbp7 | 636.70 | 3.6271 | 0.6040 | 0.0003 | 0.0147 |
| ENSMUSG00000036438 | Calm2 | 1736.06 | -3.3599 | -0.3195 | 0.0008 | 0.0282 |

| | | | | | | |
|---|---|---|---|---|---|---|
| ENSMUSG00000036469 | 1-Mar | 20.42 | 3.5178 | 2.3494 | 0.0004 | 0.0191 |
| ENSMUSG00000036504 | Phpt1 | 434.03 | -3.1777 | -0.3767 | 0.0015 | 0.0419 |
| ENSMUSG00000036528 | Ppfibp2 | 323.16 | -3.7142 | -0.6246 | 0.0002 | 0.0120 |
| ENSMUSG00000036533 | Cdc42ep3 | 470.67 | -3.2361 | -0.4678 | 0.0012 | 0.0370 |
| ENSMUSG00000036550 | Cnot1 | 1600.71 | 3.3876 | 0.2948 | 0.0007 | 0.0268 |
| ENSMUSG00000036570 | Fxyd1 | 1941.27 | -3.3938 | -0.3523 | 0.0007 | 0.0265 |
| ENSMUSG00000036606 | Plxnb2 | 229.91 | 3.3682 | 0.7068 | 0.0008 | 0.0280 |
| ENSMUSG00000036698 | Ago2 | 2258.47 | 3.5116 | 0.2583 | 0.0004 | 0.0195 |
| ENSMUSG00000036748 | Cuedc2 | 368.04 | -3.2420 | -0.3364 | 0.0012 | 0.0369 |
| ENSMUSG00000036751 | Cox6b1 | 3019.16 | -3.3358 | -0.3284 | 0.0009 | 0.0298 |
| ENSMUSG00000036887 | C1qa | 248.62 | 3.2014 | 1.5788 | 0.0014 | 0.0400 |
| ENSMUSG00000036896 | C1qc | 191.60 | 3.4049 | 1.7284 | 0.0007 | 0.0260 |
| ENSMUSG00000036902 | Neto2 | 54.29 | -3.6107 | -1.0687 | 0.0003 | 0.0153 |
| ENSMUSG00000036905 | C1qb | 153.30 | 4.6961 | 1.7780 | 0.0000 | 0.0006 |
| ENSMUSG00000037095 | Lrg1 | 218.93 | 6.1822 | 1.7129 | 0.0000 | 0.0000 |
| ENSMUSG00000037112 | Sik2 | 335.52 | 3.5400 | 0.5554 | 0.0004 | 0.0182 |
| ENSMUSG00000037152 | Ndufc1 | 1067.22 | -3.6196 | -0.4272 | 0.0003 | 0.0150 |
| ENSMUSG00000037225 | Fgf2 | 400.92 | 3.6084 | 0.6069 | 0.0003 | 0.0154 |
| ENSMUSG00000037286 | Stag1 | 602.20 | 3.1764 | 0.2948 | 0.0015 | 0.0419 |
| ENSMUSG00000037287 | Tbcel | 411.92 | -3.1617 | -0.3757 | 0.0016 | 0.0430 |
| ENSMUSG00000037405 | Icam1 | 88.71 | 3.8323 | 0.7972 | 0.0001 | 0.0087 |
| ENSMUSG00000037434 | Slc30a1 | 240.78 | 3.6787 | 0.6979 | 0.0002 | 0.0129 |
| ENSMUSG00000037487 | Ubr5 | 2870.71 | 4.0846 | 0.2944 | 0.0000 | 0.0042 |
| ENSMUSG00000037709 | Fam13a | 330.72 | 3.0937 | 0.5700 | 0.0020 | 0.0496 |
| ENSMUSG00000037736 | Limch1 | 3202.20 | 8.0694 | 0.5241 | 0.0000 | 0.0000 |
| ENSMUSG00000037820 | Tgm2 | 1084.68 | 4.5383 | 0.5479 | 0.0000 | 0.0011 |
| ENSMUSG00000037872 | Ackr1 | 77.58 | 5.6913 | 1.3570 | 0.0000 | 0.0000 |
| ENSMUSG00000037887 | Dusp8 | 418.54 | -3.9308 | -0.7629 | 0.0001 | 0.0068 |
| ENSMUSG00000037916 | Ndufv1 | 3475.74 | -3.1717 | -0.3268 | 0.0015 | 0.0423 |
| ENSMUSG00000037936 | Scarb1 | 497.06 | 5.3735 | 1.2210 | 0.0000 | 0.0000 |
| ENSMUSG00000038007 | Acer2 | 108.31 | 3.5413 | 1.3910 | 0.0004 | 0.0182 |
| ENSMUSG00000038147 | Cd84 | 22.05 | 3.4184 | 1.7788 | 0.0006 | 0.0251 |
| ENSMUSG00000038195 | Rilp | 292.71 | -3.5956 | -0.5646 | 0.0003 | 0.0160 |
| ENSMUSG00000038462 | Uqcrfs1 | 4050.17 | -3.3380 | -0.3610 | 0.0008 | 0.0298 |
| ENSMUSG00000038594 | Cep85l | 999.52 | 4.0970 | 1.6874 | 0.0000 | 0.0041 |
| ENSMUSG00000038612 | Mcl1 | 2428.34 | 3.7424 | 0.4833 | 0.0002 | 0.0111 |
| ENSMUSG00000038642 | Ctss | 111.07 | 4.1378 | 1.6009 | 0.0000 | 0.0036 |
| ENSMUSG00000038690 | Atp5j2 | 2833.02 | -3.9438 | -0.4209 | 0.0001 | 0.0065 |
| ENSMUSG00000038764 | Ptpn3 | 1952.40 | 3.2724 | 0.6426 | 0.0011 | 0.0347 |
| ENSMUSG00000038845 | Phb | 1459.22 | -3.3192 | -0.2909 | 0.0009 | 0.0310 |
| ENSMUSG00000039037 | St6galnac5 | 26.27 | 3.1767 | 1.6651 | 0.0015 | 0.0419 |
| ENSMUSG00000039105 | Atp6v1g1 | 3630.63 | -3.3554 | -0.3398 | 0.0008 | 0.0285 |
| ENSMUSG00000039109 | F13a1 | 422.53 | 4.4764 | 2.0519 | 0.0000 | 0.0013 |
| ENSMUSG00000039286 | Fndc3b | 303.50 | 4.1946 | 0.5457 | 0.0000 | 0.0033 |
| ENSMUSG00000039529 | Atp8b1 | 121.41 | 4.5857 | 1.2541 | 0.0000 | 0.0009 |
| ENSMUSG00000039585 | Myo9a | 423.81 | 4.0164 | 0.4481 | 0.0001 | 0.0053 |
| ENSMUSG00000039704 | Lmbrd2 | 599.18 | 3.2269 | 0.3092 | 0.0013 | 0.0374 |
| ENSMUSG00000039747 | Orai2 | 19.13 | 3.2785 | 1.6746 | 0.0010 | 0.0341 |

| | | | | | | |
|---|---|---|---|---|---|---|
| ENSMUSG00000039782 | Cpeb2 | 688.25 | -3.2931 | -0.5019 | 0.0010 | 0.0328 |
| ENSMUSG00000039914 | Coq10a | 2451.00 | -4.0664 | -0.4049 | 0.0000 | 0.0044 |
| ENSMUSG00000039959 | Hip1 | 616.11 | 4.5225 | 0.4825 | 0.0000 | 0.0011 |
| ENSMUSG00000039987 | Phtf2 | 4103.37 | 4.4376 | 0.3079 | 0.0000 | 0.0015 |
| ENSMUSG00000040113 | Mettl11b | 272.40 | -3.1522 | -0.5517 | 0.0016 | 0.0438 |
| ENSMUSG00000040147 | Maob | 427.07 | -3.4404 | -0.4755 | 0.0006 | 0.0238 |
| ENSMUSG00000040170 | Fmo2 | 516.15 | 4.1200 | 0.7860 | 0.0000 | 0.0038 |
| ENSMUSG00000040249 | Lrp1 | 1692.93 | 3.3279 | 0.7475 | 0.0009 | 0.0303 |
| ENSMUSG00000040269 | Mrps28 | 313.15 | -3.3658 | -0.4494 | 0.0008 | 0.0281 |
| ENSMUSG00000040407 | Akap9 | 2384.90 | 3.3601 | 0.2398 | 0.0008 | 0.0282 |
| ENSMUSG00000040485 | Lrrc52 | 35.40 | -3.7494 | -1.4677 | 0.0002 | 0.0109 |
| ENSMUSG00000040522 | Tlr8 | 10.02 | 3.1157 | 2.0687 | 0.0018 | 0.0472 |
| ENSMUSG00000040543 | Pitpnm3 | 94.02 | -3.9304 | -0.6995 | 0.0001 | 0.0068 |
| ENSMUSG00000040552 | C3ar1 | 46.70 | 3.7629 | 1.6205 | 0.0002 | 0.0104 |
| ENSMUSG00000040613 | Apobec1 | 29.68 | 3.2382 | 1.4742 | 0.0012 | 0.0370 |
| ENSMUSG00000040705 | A930016O22Rik | 196.71 | -4.9169 | -0.6847 | 0.0000 | 0.0003 |
| ENSMUSG00000040747 | Cd53 | 40.10 | 3.4286 | 1.7292 | 0.0006 | 0.0244 |
| ENSMUSG00000040820 | Hlcs | 227.28 | 4.1349 | 0.5130 | 0.0000 | 0.0037 |
| ENSMUSG00000041126 | H2afv | 446.12 | -3.0903 | -0.3592 | 0.0020 | 0.0499 |
| ENSMUSG00000041193 | Pla2g5 | 72.97 | -4.0700 | -1.7142 | 0.0000 | 0.0044 |
| ENSMUSG00000041479 | Syt15 | 66.89 | 4.7863 | 1.4294 | 0.0000 | 0.0004 |
| ENSMUSG00000041650 | Pcca | 529.56 | -3.3905 | -0.3185 | 0.0007 | 0.0267 |
| ENSMUSG00000042045 | Sln | 75.24 | 3.2073 | 1.4798 | 0.0013 | 0.0393 |
| ENSMUSG00000042784 | Muc1 | 18.12 | 3.6212 | 2.3376 | 0.0003 | 0.0149 |
| ENSMUSG00000042828 | Trim72 | 4001.78 | 10.347 | 0.9090 | 0.0000 | 0.0000 |
| ENSMUSG00000043122 | A530016L24Rik | 35.64 | 3.3409 | 1.8558 | 0.0008 | 0.0297 |
| ENSMUSG00000043535 | Setx | 1036.06 | 3.5245 | 0.3516 | 0.0004 | 0.0188 |
| ENSMUSG00000043644 | 0610009L18Rik | 97.63 | -3.1522 | -0.5861 | 0.0016 | 0.0438 |
| ENSMUSG00000043909 | Trp53bp1 | 371.94 | 3.5419 | 0.4437 | 0.0004 | 0.0182 |
| ENSMUSG00000043940 | Wdfy3 | 925.54 | 3.8903 | 0.4083 | 0.0001 | 0.0076 |
| ENSMUSG00000044308 | Ubr3 | 8051.57 | 3.2549 | 0.3947 | 0.0011 | 0.0359 |
| ENSMUSG00000044583 | Tlr7 | 21.66 | 3.4010 | 1.3528 | 0.0007 | 0.0261 |
| ENSMUSG00000044734 | Serpinb1a | 124.01 | 5.3280 | 1.7971 | 0.0000 | 0.0000 |
| ENSMUSG00000044786 | Zfp36 | 212.61 | 4.1822 | 1.2700 | 0.0000 | 0.0033 |
| ENSMUSG00000044894 | Uqcrq | 2201.83 | -3.8630 | -0.4126 | 0.0001 | 0.0081 |
| ENSMUSG00000045036 | Tmem232 | 259.34 | -3.7626 | -0.8099 | 0.0002 | 0.0104 |
| ENSMUSG00000045103 | Dmd | 3862.47 | 3.1612 | 0.5194 | 0.0016 | 0.0430 |
| ENSMUSG00000045545 | Krt14 | 54.45 | 3.1547 | 6.1571 | 0.0016 | 0.0437 |
| ENSMUSG00000045761 | Togaram2 | 66.67 | 3.9263 | 1.7118 | 0.0001 | 0.0068 |
| ENSMUSG00000045776 | Lrtm1 | 902.31 | -4.7150 | -1.0593 | 0.0000 | 0.0006 |
| ENSMUSG00000045817 | Zfp36l2 | 349.37 | 4.0807 | 0.7574 | 0.0000 | 0.0043 |
| ENSMUSG00000046532 | Ar | 1164.86 | -3.3717 | -0.4428 | 0.0007 | 0.0278 |
| ENSMUSG00000046805 | Mpeg1 | 46.42 | 3.1535 | 1.3101 | 0.0016 | 0.0438 |
| ENSMUSG00000047246 | Hist1h2be | 228.36 | -3.1351 | -0.4853 | 0.0017 | 0.0455 |
| ENSMUSG00000047342 | Zfp286 | 17.79 | -3.2519 | -1.3382 | 0.0011 | 0.0360 |
| ENSMUSG00000047592 | Nxpe5 | 46.90 | 4.6526 | 2.5224 | 0.0000 | 0.0007 |
| ENSMUSG00000047731 | Wbp1l | 1167.66 | -3.8409 | -0.3239 | 0.0001 | 0.0085 |
| ENSMUSG00000047747 | Rnf150 | 1452.81 | 3.1676 | 0.4596 | 0.0015 | 0.0426 |

| | | | | | | |
|---|---|---|---|---|---|---|
| ENSMUSG00000047793 | Sned1 | 524.12 | 5.5958 | 1.2290 | 0.0000 | 0.0000 |
| ENSMUSG00000048040 | Arxes2 | 30.10 | 3.7800 | 2.6293 | 0.0002 | 0.0099 |
| ENSMUSG00000048120 | Entpd1 | 325.29 | 4.3756 | 0.7019 | 0.0000 | 0.0019 |
| ENSMUSG00000048490 | Nrip1 | 909.26 | 4.8340 | 1.1429 | 0.0000 | 0.0004 |
| ENSMUSG00000048534 | Jaml | 15.47 | 3.4147 | 1.9310 | 0.0006 | 0.0253 |
| ENSMUSG00000048572 | Tmem252 | 19.65 | 3.9530 | 2.1534 | 0.0001 | 0.0064 |
| ENSMUSG00000048616 | Nog | 137.52 | 3.4313 | 1.1323 | 0.0006 | 0.0242 |
| ENSMUSG00000048731 | Ggnbp1 | 203.51 | -3.6337 | -0.6839 | 0.0003 | 0.0144 |
| ENSMUSG00000048865 | Arhgap30 | 43.07 | 3.4708 | 1.4364 | 0.0005 | 0.0219 |
| ENSMUSG00000048960 | Prex2 | 301.75 | 3.8708 | 0.6562 | 0.0001 | 0.0080 |
| ENSMUSG00000049076 | Acap2 | 286.24 | 3.3934 | 0.4300 | 0.0007 | 0.0265 |
| ENSMUSG00000049103 | Ccr2 | 38.84 | 4.1447 | 2.1431 | 0.0000 | 0.0036 |
| ENSMUSG00000049130 | C5ar1 | 54.81 | 3.5282 | 1.6865 | 0.0004 | 0.0187 |
| ENSMUSG00000049281 | Scn3b | 68.70 | 4.6863 | 1.8742 | 0.0000 | 0.0006 |
| ENSMUSG00000049422 | Chchd10 | 3527.04 | -3.8655 | -0.4066 | 0.0001 | 0.0081 |
| ENSMUSG00000049791 | Fzd4 | 1571.64 | 3.2378 | 1.0617 | 0.0012 | 0.0370 |
| ENSMUSG00000049892 | Rasd1 | 28.29 | 4.0942 | 2.0319 | 0.0000 | 0.0041 |
| ENSMUSG00000050105 | Grrp1 | 49.44 | 4.1631 | 1.2033 | 0.0000 | 0.0034 |
| ENSMUSG00000050122 | Vwa3b | 65.99 | -3.1224 | -0.9894 | 0.0018 | 0.0467 |
| ENSMUSG00000050821 | Fam131a | 423.87 | -3.0960 | -0.6040 | 0.0020 | 0.0494 |
| ENSMUSG00000050931 | Sgms2 | 35.06 | 3.1267 | 1.0601 | 0.0018 | 0.0461 |
| ENSMUSG00000051344 | Plekhm3 | 183.68 | 3.2302 | 0.4469 | 0.0012 | 0.0371 |
| ENSMUSG00000051439 | Cd14 | 37.40 | 3.6993 | 1.5734 | 0.0002 | 0.0125 |
| ENSMUSG00000052033 | Pfdn4 | 92.10 | -3.2467 | -0.5796 | 0.0012 | 0.0365 |
| ENSMUSG00000052276 | Ostn | 960.57 | -4.3716 | -2.0667 | 0.0000 | 0.0019 |
| ENSMUSG00000052299 | Ltn1 | 725.92 | 3.0926 | 0.2682 | 0.0020 | 0.0497 |
| ENSMUSG00000052565 | Hist1h1d | 115.87 | -3.1914 | -0.5920 | 0.0014 | 0.0407 |
| ENSMUSG00000052738 | Suclg1 | 2450.95 | -5.4345 | -0.4467 | 0.0000 | 0.0000 |
| ENSMUSG00000052821 | Cysltr1 | 46.20 | 4.1723 | 1.3932 | 0.0000 | 0.0034 |
| ENSMUSG00000052837 | Junb | 220.46 | 5.3275 | 1.6523 | 0.0000 | 0.0000 |
| ENSMUSG00000052920 | Prkg1 | 851.30 | 3.3014 | 0.5509 | 0.0010 | 0.0324 |
| ENSMUSG00000053113 | Socs3 | 251.37 | 6.0678 | 2.8560 | 0.0000 | 0.0000 |
| ENSMUSG00000053175 | Bcl3 | 30.64 | 3.7716 | 1.5787 | 0.0002 | 0.0101 |
| ENSMUSG00000053453 | Thoc7 | 727.26 | -3.8207 | -0.3954 | 0.0001 | 0.0089 |
| ENSMUSG00000053626 | Tll1 | 29.73 | 3.6655 | 1.3147 | 0.0002 | 0.0132 |
| ENSMUSG00000053716 | Dusp7 | 282.42 | 3.1365 | 0.6917 | 0.0017 | 0.0454 |
| ENSMUSG00000053801 | Grwd1 | 125.03 | -3.0915 | -0.5163 | 0.0020 | 0.0498 |
| ENSMUSG00000054312 | Mrps21 | 740.50 | -3.5593 | -0.3324 | 0.0004 | 0.0176 |
| ENSMUSG00000054408 | Spcs3 | 625.59 | 3.8478 | 0.5314 | 0.0001 | 0.0084 |
| ENSMUSG00000054555 | Adam12 | 19.98 | 3.8960 | 2.3736 | 0.0001 | 0.0075 |
| ENSMUSG00000054889 | Dsp | 25.80 | 3.1739 | 6.3289 | 0.0015 | 0.0421 |
| ENSMUSG00000055489 | Ano5 | 3739.35 | 5.2919 | 0.5514 | 0.0000 | 0.0001 |
| ENSMUSG00000055762 | Eef1d | 988.27 | -3.3373 | -0.2752 | 0.0008 | 0.0298 |
| ENSMUSG00000055839 | Elob | 939.75 | -3.9911 | -0.4718 | 0.0001 | 0.0056 |
| ENSMUSG00000055980 | Irs1 | 2221.15 | -3.3980 | -0.6604 | 0.0007 | 0.0263 |
| ENSMUSG00000056025 | Clca3a1 | 28.24 | 4.7378 | 2.7236 | 0.0000 | 0.0005 |
| ENSMUSG00000056608 | Chd9 | 657.38 | 4.1593 | 0.4153 | 0.0000 | 0.0034 |
| ENSMUSG00000056973 | Ces1d | 669.87 | -4.1907 | -0.7115 | 0.0000 | 0.0033 |

| | | | | | | |
|---|---|---|---|---|---|---|
| ENSMUSG00000057719 | Sh3rf2 | 1019.84 | 3.4426 | 0.6254 | 0.0006 | 0.0237 |
| ENSMUSG00000058076 | Sdhc | 3451.54 | -3.1807 | -0.3261 | 0.0015 | 0.0416 |
| ENSMUSG00000058239 | Usf2 | 717.57 | -3.0921 | -0.2862 | 0.0020 | 0.0497 |
| ENSMUSG00000058624 | Gda | 392.23 | 3.2690 | 1.2674 | 0.0011 | 0.0348 |
| ENSMUSG00000058818 | Pirb | 62.97 | 3.0962 | 1.6414 | 0.0020 | 0.0494 |
| ENSMUSG00000058927 | ENSMUSG00000058927 | 4573.70 | -4.0134 | -0.4487 | 0.0001 | 0.0053 |
| ENSMUSG00000058966 | Fam57b | 951.85 | -3.3626 | -0.5179 | 0.0008 | 0.0282 |
| ENSMUSG00000059182 | Skap2 | 107.97 | 3.4965 | 0.7146 | 0.0005 | 0.0203 |
| ENSMUSG00000059534 | Uqcr10 | 1876.59 | -3.2870 | -0.4134 | 0.0010 | 0.0332 |
| ENSMUSG00000059866 | Tnip2 | 101.02 | 3.5186 | 0.7633 | 0.0004 | 0.0191 |
| ENSMUSG00000059890 | Ube4a | 974.41 | 3.6074 | 0.4377 | 0.0003 | 0.0154 |
| ENSMUSG00000059970 | Hspa2 | 48.32 | 3.2214 | 0.8350 | 0.0013 | 0.0379 |
| ENSMUSG00000060063 | Alox5ap | 38.51 | 3.3616 | 1.2635 | 0.0008 | 0.0282 |
| ENSMUSG00000060376 | Bckdha | 1563.79 | -3.9566 | -0.4779 | 0.0001 | 0.0063 |
| ENSMUSG00000060586 | H2-Eb1 | 77.66 | 3.1291 | 1.1632 | 0.0018 | 0.0460 |
| ENSMUSG00000060591 | Ifitm2 | 344.66 | 3.1621 | 0.5913 | 0.0016 | 0.0430 |
| ENSMUSG00000061518 | Cox5b | 3195.68 | -3.8550 | -0.3781 | 0.0001 | 0.0083 |
| ENSMUSG00000061723 | Tnnt3 | 194619.51 | -3.2511 | -0.3359 | 0.0011 | 0.0360 |
| ENSMUSG00000061758 | Akr1b10 | 425.59 | -3.8257 | -0.6203 | 0.0001 | 0.0088 |
| ENSMUSG00000061838 | Suclg2 | 718.17 | -3.5010 | -0.3857 | 0.0005 | 0.0200 |
| ENSMUSG00000062593 | Lilrb4a | 7.68 | 3.8735 | 5.7569 | 0.0001 | 0.0079 |
| ENSMUSG00000062609 | Kcnj15 | 39.30 | 3.4353 | 1.7622 | 0.0006 | 0.0240 |
| ENSMUSG00000062960 | Kdr | 747.10 | -4.1325 | -0.5827 | 0.0000 | 0.0037 |
| ENSMUSG00000062981 | Mrpl42 | 1720.01 | -3.1143 | -0.3443 | 0.0018 | 0.0474 |
| ENSMUSG00000063236 | 1110038F14Rik | 117.03 | -3.6434 | -0.5722 | 0.0003 | 0.0140 |
| ENSMUSG00000063564 | Col23a1 | 171.32 | 3.5563 | 0.8444 | 0.0004 | 0.0177 |
| ENSMUSG00000063694 | Cycs | 2402.94 | -3.1892 | -0.3695 | 0.0014 | 0.0408 |
| ENSMUSG00000064057 | Scgb3a1 | 5.94 | 3.8751 | 5.8339 | 0.0001 | 0.0079 |
| ENSMUSG00000064339 | ENSMUSG00000064339 | 49475.09 | -3.6891 | -0.5564 | 0.0002 | 0.0127 |
| ENSMUSG00000064341 | ND1 | 153401.59 | -5.1056 | -0.5178 | 0.0000 | 0.0001 |
| ENSMUSG00000064345 | ND2 | 133107.24 | -4.0764 | -0.4617 | 0.0000 | 0.0043 |
| ENSMUSG00000064351 | COX1 | 514862.61 | -3.2286 | -0.3677 | 0.0012 | 0.0373 |
| ENSMUSG00000064356 | ATP8 | 24804.79 | -3.2223 | -0.3734 | 0.0013 | 0.0379 |
| ENSMUSG00000064357 | ATP6 | 18914.57 | -3.6401 | -0.5121 | 0.0003 | 0.0141 |
| ENSMUSG00000064358 | COX3 | 43118.87 | -4.6147 | -0.8789 | 0.0000 | 0.0008 |
| ENSMUSG00000064360 | ND3 | 7440.53 | -4.0372 | -0.5844 | 0.0001 | 0.0049 |
| ENSMUSG00000064363 | ND4 | 195059.13 | -4.4392 | -0.4560 | 0.0000 | 0.0015 |
| ENSMUSG00000064368 | ND6 | 9064.28 | -4.5373 | -0.5500 | 0.0000 | 0.0011 |
| ENSMUSG00000064370 | CYTB | 189105.17 | -4.1072 | -0.4496 | 0.0000 | 0.0040 |
| ENSMUSG00000065947 | ND4L | 13944.73 | -3.9691 | -0.4215 | 0.0001 | 0.0061 |
| ENSMUSG00000066800 | Rnasel | 70.02 | 3.2113 | 1.0815 | 0.0013 | 0.0390 |
| ENSMUSG00000066839 | Ecsit | 865.53 | -4.6793 | -0.4095 | 0.0000 | 0.0006 |
| ENSMUSG00000068114 | Ccdc134 | 103.70 | -3.3278 | -0.6581 | 0.0009 | 0.0303 |
| ENSMUSG00000068697 | Myoz1 | 18976.94 | -3.1916 | -0.2612 | 0.0014 | 0.0407 |
| ENSMUSG00000069516 | Lyz2 | 663.57 | 3.1296 | 1.3012 | 0.0018 | 0.0460 |
| ENSMUSG00000069806 | Cacng7 | 91.53 | -4.0005 | -0.7960 | 0.0001 | 0.0055 |
| ENSMUSG00000069833 | Ahnak | 11268.76 | 3.6170 | 0.3921 | 0.0003 | 0.0150 |
| ENSMUSG00000070424 | Art5 | 339.31 | 4.3442 | 0.5923 | 0.0000 | 0.0020 |

| | | | | | | |
|---|---|---|---|---|---|---|
| ENSMUSG00000070498 | Tmem132b | 51.32 | -3.5595 | -0.9805 | 0.0004 | 0.0176 |
| ENSMUSG00000071005 | Ccl19 | 99.72 | -3.8765 | -0.9616 | 0.0001 | 0.0079 |
| ENSMUSG00000071014 | Ndufb6 | 1021.15 | -3.1845 | -0.2975 | 0.0015 | 0.0413 |
| ENSMUSG00000071528 | Usmg5 | 1802.29 | -3.1441 | -0.3852 | 0.0017 | 0.0447 |
| ENSMUSG00000071637 | Cebpd | 153.84 | 4.3943 | 1.5741 | 0.0000 | 0.0017 |
| ENSMUSG00000071711 | Mpst | 451.00 | -3.1839 | -0.3988 | 0.0015 | 0.0413 |
| ENSMUSG00000071713 | Csf2rb | 168.55 | 7.3619 | 2.1959 | 0.0000 | 0.0000 |
| ENSMUSG00000071714 | Csf2rb2 | 124.50 | 6.6344 | 2.1159 | 0.0000 | 0.0000 |
| ENSMUSG00000073016 | Uprt | 96.29 | 3.1647 | 0.7418 | 0.0016 | 0.0428 |
| ENSMUSG00000073418 | C4b | 1231.64 | 3.6776 | 2.5384 | 0.0002 | 0.0129 |
| ENSMUSG00000073421 | H2-Ab1 | 65.82 | 3.1158 | 1.2331 | 0.0018 | 0.0472 |
| ENSMUSG00000073489 | Ifi204 | 81.01 | 4.6292 | 1.1320 | 0.0000 | 0.0008 |
| ENSMUSG00000073490 | Ifi207 | 96.12 | 3.9296 | 1.6848 | 0.0001 | 0.0068 |
| ENSMUSG00000074207 | Adh1 | 141.51 | 3.3947 | 0.8162 | 0.0007 | 0.0265 |
| ENSMUSG00000074218 | Cox7a1 | 1242.93 | -4.4404 | -0.6255 | 0.0000 | 0.0015 |
| ENSMUSG00000074417 | Pira11\|Pira7\|Gm14548 | 9.07 | 3.2329 | 3.2035 | 0.0012 | 0.0370 |
| ENSMUSG00000074794 | Arrdc3 | 661.73 | 3.5900 | 0.6612 | 0.0003 | 0.0161 |
| ENSMUSG00000075467 | Dnlz | 321.43 | -3.2323 | -0.4101 | 0.0012 | 0.0370 |
| ENSMUSG00000075602 | Ly6a | 711.60 | 4.0078 | 0.7476 | 0.0001 | 0.0054 |
| ENSMUSG00000078673 | Gm2083\|Mup19 | 28.17 | 4.2456 | 8.3693 | 0.0000 | 0.0028 |
| ENSMUSG00000079168 | Cd209g | 24.17 | 3.0994 | 1.8718 | 0.0019 | 0.0490 |
| ENSMUSG00000079173 | Zan | 18.14 | 4.8535 | 4.3147 | 0.0000 | 0.0004 |
| ENSMUSG00000079227 | Ccr5 | 19.14 | 3.2362 | 2.4199 | 0.0012 | 0.0370 |
| ENSMUSG00000079419 | Ms4a6c | 40.20 | 4.1642 | 1.6193 | 0.0000 | 0.0034 |
| ENSMUSG00000079508 | Apoo | 1359.64 | -3.5799 | -0.3320 | 0.0003 | 0.0165 |
| ENSMUSG00000080893 | ENSMUSG00000080893 | 304.15 | -4.0700 | -0.5687 | 0.0000 | 0.0044 |
| ENSMUSG00000081752 | ENSMUSG00000081752 | 20.82 | 5.4261 | 19.9693 | 0.0000 | 0.0000 |
| ENSMUSG00000083626 | ENSMUSG00000083626 | 157.00 | -3.5165 | -0.6430 | 0.0004 | 0.0192 |
| ENSMUSG00000084843 | ENSMUSG00000084843 | 38.50 | -3.5245 | -1.4525 | 0.0004 | 0.0188 |
| ENSMUSG00000085133 | ENSMUSG00000085133 | 44.96 | 3.4866 | 1.1612 | 0.0005 | 0.0209 |
| ENSMUSG00000086866 | ENSMUSG00000086866 | 18.94 | 4.5859 | 3.1026 | 0.0000 | 0.0009 |
| ENSMUSG00000089704 | Galnt2 | 649.65 | 4.2317 | 0.6320 | 0.0000 | 0.0029 |
| ENSMUSG00000089719 | ENSMUSG00000089719 | 10.72 | 3.2880 | 2.4207 | 0.0010 | 0.0332 |
| ENSMUSG00000090258 | Churc1 | 339.84 | -3.8647 | -0.4629 | 0.0001 | 0.0081 |
| ENSMUSG00000091712 | Sec14l5 | 697.75 | -3.2400 | -0.4547 | 0.0012 | 0.0370 |
| ENSMUSG00000094720 | n-R5s100 | 53.67 | -3.4357 | -2.9070 | 0.0006 | 0.0240 |
| ENSMUSG00000095298 | ENSMUSG00000095298 | 12.14 | -3.2408 | -1.8752 | 0.0012 | 0.0370 |
| ENSMUSG00000096146 | Kcnj11 | 2350.05 | -3.2301 | -0.3330 | 0.0012 | 0.0371 |
| ENSMUSG00000096793 | ENSMUSG00000096793 | 10.01 | -3.4461 | -2.3246 | 0.0006 | 0.0235 |
| ENSMUSG00000097487 | Ptges3l | 467.86 | -3.2389 | -0.5637 | 0.0012 | 0.0370 |
| ENSMUSG00000099021 | Rn7s1 | 1277.45 | -5.1734 | -1.1868 | 0.0000 | 0.0001 |
| ENSMUSG00000100550 | 2310039L15Rik | 17.81 | -3.5068 | -1.9487 | 0.0005 | 0.0197 |
| ENSMUSG00000100862 | ENSMUSG00000100862 | 146249.19 | -5.0028 | -0.5269 | 0.0000 | 0.0002 |
| ENSMUSG00000101111 | ENSMUSG00000101111 | 153152.83 | -3.8491 | -0.4910 | 0.0001 | 0.0084 |
| ENSMUSG00000101249 | ENSMUSG00000101249 | 16141.86 | -10.634 | -1.6575 | 0.0000 | 0.0000 |
| ENSMUSG00000101655 | 2310040G24Rik | 625.36 | -3.9564 | -0.5603 | 0.0001 | 0.0063 |
| ENSMUSG00000101939 | ENSMUSG00000101939 | 14756.84 | -3.8343 | -0.5937 | 0.0001 | 0.0087 |
| ENSMUSG00000102070 | ENSMUSG00000102070 | 159101.20 | -4.9984 | -0.5538 | 0.0000 | 0.0002 |

| | | | | | | |
|---|---|---|---|---|---|---|
| ENSMUSG00000102573 | ENSMUSG00000102573 | 194.15 | 3.8306 | 0.6782 | 0.0001 | 0.0087 |
| ENSMUSG00000103560 | ENSMUSG00000103560 | 15.00 | 3.6888 | 2.3397 | 0.0002 | 0.0127 |
| ENSMUSG00000104222 | ENSMUSG00000104222 | 1187.58 | 3.3870 | 0.4235 | 0.0007 | 0.0268 |
| ENSMUSG00000104453 | ENSMUSG00000104453 | 1670.69 | -3.1038 | -0.3890 | 0.0019 | 0.0485 |
| ENSMUSG00000106840 | ENSMUSG00000106840 | 114.40 | -3.1393 | -0.6395 | 0.0017 | 0.0451 |
| ENSMUSG00000106892 | ENSMUSG00000106892 | 5.43 | 3.8455 | 5.6833 | 0.0001 | 0.0085 |
| ENSMUSG00000107689 | ENSMUSG00000107689 | 281.53 | -3.8942 | -0.5992 | 0.0001 | 0.0075 |
| ENSMUSG00000107881 | ENSMUSG00000107881 | 757.66 | -4.0121 | -0.3909 | 0.0001 | 0.0053 |
| ENSMUSG00000109909 | ENSMUSG00000109909 | 82.80 | 3.1669 | 0.6457 | 0.0015 | 0.0426 |
| ENSMUSG00000110588 | ENSMUSG00000110588 | 16.02 | 4.1414 | 4.2713 | 0.0000 | 0.0036 |
| ENSMUSG00000110779 | ENSMUSG00000110779 | 511.02 | 3.8361 | 0.7932 | 0.0001 | 0.0087 |
| ENSMUSG00000111942 | ENSMUSG00000111942 | 6.69 | 3.2440 | 4.6005 | 0.0012 | 0.0368 |
| ENSMUSG00000112182 | ENSMUSG00000112182 | 23.02 | -3.1821 | -1.1001 | 0.0015 | 0.0415 |
| ENSMUSG00000113780 | ENSMUSG00000113780 | 18.65 | -3.9463 | -1.6540 | 0.0001 | 0.0065 |
| ENSMUSG00000113902 | ENSMUSG00000113902 | 939.79 | -3.4325 | -0.4716 | 0.0006 | 0.0242 |
| ENSMUSG00000114019 | ENSMUSG00000114019 | 249.45 | -4.4872 | -0.5508 | 0.0000 | 0.0013 |
| ENSMUSG00000114608 | ENSMUSG00000114608 | 21.25 | 3.5527 | 1.8184 | 0.0004 | 0.0178 |
| ENSMUSG00000116066 | ENSMUSG00000116066 | 176.29 | -3.1592 | -0.7933 | 0.0016 | 0.0432 |
| ENSMUSG00000118167 | ENSMUSG00000118167 | 6.13 | -3.9251 | -4.5074 | 0.0001 | 0.0068 |
| ENSMUSG00000118206 | ENSMUSG00000118206 | 103.63 | 3.4694 | 1.1459 | 0.0005 | 0.0219 |
