## Supplemental Table 3 for "Pioglitazone administration restores a PPARG-dependent transcriptional network and ATP levels within skeletal muscles of mice implanted with patient-derived breast tumors"

**Supplemental Table 3: Differentially expressed genes in muscles of BC-PDOX PIO mice compared to NSG-Con.**

| GeneID | Gene | baseMean | stat | log2FoldChange | pvalue | padj |
|---|---|---|---|---|---|---|
| ENSMUSG00000000305 | Cdh4 | 371.62 | -5.16552 | -1.56350 | 0.00000 | 0.00024 |
| ENSMUSG00000001175 | Calm1 | 10367.67 | -4.26128 | -0.40075 | 0.00002 | 0.00588 |
| ENSMUSG00000002107 | Celf2 | 2403.92 | -3.74383 | -0.29223 | 0.00018 | 0.02334 |
| ENSMUSG00000003226 | Ranbp2 | 1668.60 | 3.72483 | 0.39905 | 0.00020 | 0.02425 |
| ENSMUSG00000004040 | Stat3 | 1781.13 | 9.96860 | 1.18959 | 0.00000 | 0.00000 |
| ENSMUSG00000004098 | Col5a3 | 509.84 | -4.19248 | -0.88298 | 0.00003 | 0.00690 |
| ENSMUSG00000004939 | Nmrk2 | 511.01 | -3.82484 | -1.80656 | 0.00013 | 0.01987 |
| ENSMUSG00000005580 | Adcy9 | 627.19 | 3.76426 | 0.50086 | 0.00017 | 0.02196 |
| ENSMUSG00000006362 | Cbfa2t3 | 416.61 | -3.94342 | -0.53836 | 0.00008 | 0.01450 |
| ENSMUSG00000008035 | Mid1ip1 | 348.49 | 3.87061 | 1.25635 | 0.00011 | 0.01781 |
| ENSMUSG00000009185 | Ccl8 | 134.01 | 4.40114 | 4.87273 | 0.00001 | 0.00397 |
| ENSMUSG00000009376 | Met | 321.39 | 4.61911 | 0.84401 | 0.00000 | 0.00184 |
| ENSMUSG00000009647 | Mcu | 873.30 | 5.94957 | 0.53035 | 0.00000 | 0.00001 |
| ENSMUSG00000010651 | Acaa1b | 11.07 | 3.83938 | 4.59492 | 0.00012 | 0.01958 |
| ENSMUSG00000013275 | Slc41a1 | 829.19 | 4.36546 | 0.59602 | 0.00001 | 0.00436 |
| ENSMUSG00000013584 | Aldh1a2 | 100.55 | 3.98072 | 1.59103 | 0.00007 | 0.01329 |
| ENSMUSG00000014355 | Anapc1 | 803.03 | 3.51067 | 0.34605 | 0.00045 | 0.04234 |
| ENSMUSG00000014444 | Piezo1 | 893.37 | 4.66010 | 1.00681 | 0.00000 | 0.00161 |
| ENSMUSG00000015243 | Abca1 | 721.65 | 4.35233 | 1.01850 | 0.00001 | 0.00442 |
| ENSMUSG00000016756 | Cmah | 120.23 | 4.46557 | 1.08619 | 0.00001 | 0.00327 |
| ENSMUSG00000017009 | Sdc4 | 362.78 | -3.74087 | -0.65298 | 0.00018 | 0.02334 |
| ENSMUSG00000017677 | Wsb1 | 187.64 | -5.07902 | -0.91123 | 0.00000 | 0.00035 |
| ENSMUSG00000018340 | Anxa6 | 2504.20 | 4.20599 | 0.54503 | 0.00003 | 0.00662 |
| ENSMUSG00000019838 | Slc16a10 | 944.25 | -4.61861 | -0.59332 | 0.00000 | 0.00184 |
| ENSMUSG00000020017 | Hal | 122.93 | 3.79299 | 1.45334 | 0.00015 | 0.02072 |
| ENSMUSG00000020227 | Irak3 | 86.30 | 3.48976 | 0.90158 | 0.00048 | 0.04508 |
| ENSMUSG00000020354 | Sgcd | 520.07 | 4.48337 | 1.03634 | 0.00001 | 0.00310 |
| ENSMUSG00000020681 | Ace | 677.21 | 3.55315 | 0.75368 | 0.00038 | 0.03747 |
| ENSMUSG00000020719 | Ddx5 | 7066.19 | -3.70743 | -0.29618 | 0.00021 | 0.02558 |
| ENSMUSG00000020846 | Rflnb | 277.30 | -4.22277 | -0.81023 | 0.00002 | 0.00662 |
| ENSMUSG00000021200 | Asb2 | 5604.25 | -4.86098 | -0.88928 | 0.00000 | 0.00069 |
| ENSMUSG00000021288 | Klc1 | 807.72 | -3.63159 | -0.35233 | 0.00028 | 0.03036 |
| ENSMUSG00000021423 | Ly86 | 19.44 | 3.85661 | 2.39525 | 0.00011 | 0.01845 |
| ENSMUSG00000021796 | Bmpr1a | 1561.07 | 3.44938 | 0.30575 | 0.00056 | 0.04934 |
| ENSMUSG00000021815 | Mss51 | 11204.61 | -4.28378 | -0.39297 | 0.00002 | 0.00554 |
| ENSMUSG00000021950 | Anxa8 | 27.11 | 3.74297 | 2.59067 | 0.00018 | 0.02334 |
| ENSMUSG00000021983 | Atp8a2\|LOC108168164 | 116.24 | 3.82153 | 0.85890 | 0.00013 | 0.01987 |
| ENSMUSG00000022146 | Osmr | 263.68 | 4.00056 | 0.81820 | 0.00006 | 0.01261 |
| ENSMUSG00000022181 | C6 | 27.35 | 4.48457 | 4.48689 | 0.00001 | 0.00310 |
| ENSMUSG00000022895 | Ets2 | 937.01 | 4.12105 | 0.42110 | 0.00004 | 0.00884 |
| ENSMUSG00000023088 | Abcc1 | 562.39 | 3.97550 | 0.45924 | 0.00007 | 0.01329 |
| ENSMUSG00000023186 | Vwa5a | 582.70 | 4.09717 | 0.59730 | 0.00004 | 0.00934 |
| ENSMUSG00000023367 | Tmem176a | 144.96 | 4.36019 | 1.31615 | 0.00001 | 0.00436 |
| ENSMUSG00000024014 | Pim1 | 271.88 | -5.19977 | -1.30687 | 0.00000 | 0.00023 |

| | | | | | | |
|---|---|---|---|---|---|---|
| ENSMUSG00000024030 | Abcg1 | 57.96 | 3.81279 | 1.05091 | 0.00014 | 0.01987 |
| ENSMUSG00000024033 | Rsph1 | 57.71 | -5.62216 | -2.25891 | 0.00000 | 0.00003 |
| ENSMUSG00000024036 | Slc37a1 | 42.51 | -3.43693 | -1.38012 | 0.00059 | 0.04992 |
| ENSMUSG00000024589 | Nedd4l | 813.94 | 5.64540 | 0.77091 | 0.00000 | 0.00003 |
| ENSMUSG00000024597 | Slc12a2 | 1961.34 | 3.82999 | 0.47656 | 0.00013 | 0.01986 |
| ENSMUSG00000024687 | Osbp | 1691.58 | 4.45798 | 0.40365 | 0.00001 | 0.00330 |
| ENSMUSG00000024789 | Jak2 | 667.85 | 3.57710 | 0.61805 | 0.00035 | 0.03478 |
| ENSMUSG00000024935 | Slc1a1 | 63.48 | -3.72622 | -0.93327 | 0.00019 | 0.02425 |
| ENSMUSG00000025255 | Zfhx4 | 483.98 | 3.44527 | 0.37292 | 0.00057 | 0.04934 |
| ENSMUSG00000025261 | Huwe1 | 5475.64 | 3.69107 | 0.31707 | 0.00022 | 0.02647 |
| ENSMUSG00000025511 | Tspan4 | 102.67 | 3.94289 | 0.95227 | 0.00008 | 0.01450 |
| ENSMUSG00000025650 | Col7a1 | 310.95 | -3.89924 | -0.96662 | 0.00010 | 0.01657 |
| ENSMUSG00000025777 | Gdap1 | 262.69 | -4.06976 | -0.95632 | 0.00005 | 0.01007 |
| ENSMUSG00000025887 | Casp12 | 174.86 | 3.46242 | 0.79563 | 0.00054 | 0.04790 |
| ENSMUSG00000026077 | Npas2 | 26.62 | -3.46718 | -1.58717 | 0.00053 | 0.04735 |
| ENSMUSG00000026580 | Selp | 81.23 | 4.73040 | 1.43920 | 0.00000 | 0.00127 |
| ENSMUSG00000026678 | Rgs5 | 3069.12 | -3.76269 | -0.45333 | 0.00017 | 0.02196 |
| ENSMUSG00000027322 | Siglec1 | 63.41 | 3.64001 | 1.96755 | 0.00027 | 0.02982 |
| ENSMUSG00000027365 | Trpm7 | 876.79 | 3.64849 | 0.50278 | 0.00026 | 0.02906 |
| ENSMUSG00000027397 | Slc20a1 | 213.29 | -4.04013 | -0.81205 | 0.00005 | 0.01095 |
| ENSMUSG00000027533 | Fabp5 | 93.41 | 3.49789 | 0.71541 | 0.00047 | 0.04410 |
| ENSMUSG00000027546 | Atp9a | 954.53 | 6.40553 | 1.38077 | 0.00000 | 0.00000 |
| ENSMUSG00000028132 | Tmem56 | 1772.63 | 3.67648 | 0.39621 | 0.00024 | 0.02749 |
| ENSMUSG00000028156 | Eif4e | 1552.60 | 4.20663 | 0.46937 | 0.00003 | 0.00662 |
| ENSMUSG00000028195 | Cyr61 | 485.31 | 3.67367 | 0.60147 | 0.00024 | 0.02758 |
| ENSMUSG00000028197 | Col24a1 | 93.53 | -5.16513 | -1.34696 | 0.00000 | 0.00024 |
| ENSMUSG00000028525 | Pde4b | 1570.39 | -3.44408 | -0.62973 | 0.00057 | 0.04934 |
| ENSMUSG00000028600 | Podn | 181.20 | 3.65522 | 0.94777 | 0.00026 | 0.02875 |
| ENSMUSG00000028630 | Dyrk2 | 1970.17 | -4.12877 | -0.65999 | 0.00004 | 0.00875 |
| ENSMUSG00000030341 | Tnfrsf1a | 311.44 | 3.46851 | 0.74609 | 0.00052 | 0.04735 |
| ENSMUSG00000030680 | ENSMUSG00000030680 | 60.01 | -3.80673 | -9.17096 | 0.00014 | 0.01987 |
| ENSMUSG00000030737 | Slco2b1 | 208.30 | 4.01653 | 0.89482 | 0.00006 | 0.01195 |
| ENSMUSG00000030786 | Itgam | 59.27 | 3.59183 | 1.35795 | 0.00033 | 0.03390 |
| ENSMUSG00000030798 | Cd37 | 24.69 | 3.76469 | 1.92442 | 0.00017 | 0.02196 |
| ENSMUSG00000031010 | Usp9x | 4328.18 | 4.36361 | 0.36138 | 0.00001 | 0.00436 |
| ENSMUSG00000031465 | Angpt2 | 40.16 | -3.78749 | -1.16928 | 0.00015 | 0.02072 |
| ENSMUSG00000031592 | Pcm1 | 1440.55 | 4.21140 | 0.50981 | 0.00003 | 0.00662 |
| ENSMUSG00000031608 | Galnt7 | 101.49 | 3.57616 | 0.82492 | 0.00035 | 0.03478 |
| ENSMUSG00000031805 | Jak3 | 171.56 | 4.40659 | 0.73685 | 0.00001 | 0.00397 |
| ENSMUSG00000031963 | Bmper | 100.42 | 3.52896 | 0.97312 | 0.00042 | 0.04052 |
| ENSMUSG00000032013 | Trim29 | 9.16 | 3.97788 | 6.00474 | 0.00007 | 0.01329 |
| ENSMUSG00000032340 | Neo1 | 1095.32 | 3.82813 | 0.33998 | 0.00013 | 0.01986 |
| ENSMUSG00000032355 | Mlip | 1570.84 | 4.39323 | 0.39707 | 0.00001 | 0.00402 |
| ENSMUSG00000032503 | Arpp21 | 200.60 | 3.87924 | 0.97088 | 0.00010 | 0.01738 |
| ENSMUSG00000032666 | 1700025G04Rik | 1764.82 | 3.59612 | 0.29817 | 0.00032 | 0.03358 |
| ENSMUSG00000032744 | Heyl | 130.23 | -3.45997 | -0.70476 | 0.00054 | 0.04805 |
| ENSMUSG00000033082 | Clec1a | 60.93 | 3.52523 | 0.96171 | 0.00042 | 0.04083 |
| ENSMUSG00000033624 | Pdpr | 2172.20 | 3.83055 | 0.52909 | 0.00013 | 0.01986 |

| | | | | | | |
|---|---|---|---|---|---|---|
| ENSMUSG00000034021 | Pds5b | 1039.82 | 3.61310 | 0.35232 | 0.00030 | 0.03191 |
| ENSMUSG00000034040 | Galnt17 | 173.35 | 5.30847 | 1.26435 | 0.00000 | 0.00015 |
| ENSMUSG00000034593 | Myo5a | 257.43 | 3.48197 | 0.67464 | 0.00050 | 0.04593 |
| ENSMUSG00000034612 | Chst11 | 94.63 | 3.51529 | 1.34158 | 0.00044 | 0.04211 |
| ENSMUSG00000034636 | Zyg11b | 2281.73 | 4.10126 | 0.52270 | 0.00004 | 0.00933 |
| ENSMUSG00000034845 | Plvap | 380.94 | 3.48866 | 0.71815 | 0.00049 | 0.04508 |
| ENSMUSG00000034973 | Dop1a | 265.38 | 3.58068 | 0.45055 | 0.00034 | 0.03478 |
| ENSMUSG00000035284 | Vps13c | 810.22 | 3.58178 | 0.52985 | 0.00034 | 0.03478 |
| ENSMUSG00000036854 | Hspb6 | 11867.97 | 3.60392 | 0.52987 | 0.00031 | 0.03282 |
| ENSMUSG00000036905 | C1qb | 153.30 | 4.23551 | 1.70737 | 0.00002 | 0.00647 |
| ENSMUSG00000037095 | Lrg1 | 218.93 | 4.05510 | 1.20106 | 0.00005 | 0.01045 |
| ENSMUSG00000037225 | Fgf2 | 400.92 | 4.89208 | 0.87347 | 0.00000 | 0.00061 |
| ENSMUSG00000037266 | Rsrp1 | 4395.46 | -4.18577 | -0.58910 | 0.00003 | 0.00699 |
| ENSMUSG00000037736 | Limch1 | 3202.20 | 4.97969 | 0.34593 | 0.00000 | 0.00045 |
| ENSMUSG00000037872 | Ackr1 | 77.58 | 5.25641 | 1.32042 | 0.00000 | 0.00018 |
| ENSMUSG00000037887 | Dusp8 | 418.54 | -3.65480 | -0.75893 | 0.00026 | 0.02875 |
| ENSMUSG00000037936 | Scarb1 | 497.06 | 4.34163 | 1.05332 | 0.00001 | 0.00454 |
| ENSMUSG00000038007 | Acer2 | 108.31 | 3.89263 | 1.62333 | 0.00010 | 0.01683 |
| ENSMUSG00000038121 | Fam210a | 2589.97 | 3.66823 | 0.58063 | 0.00024 | 0.02774 |
| ENSMUSG00000038175 | Mylip | 445.56 | 4.29628 | 0.59788 | 0.00002 | 0.00534 |
| ENSMUSG00000038201 | Kcna7 | 834.76 | 4.21732 | 0.52500 | 0.00002 | 0.00662 |
| ENSMUSG00000038642 | Ctss | 111.07 | 3.76572 | 1.55020 | 0.00017 | 0.02196 |
| ENSMUSG00000038754 | Elovl3 | 6.13 | 3.47116 | 5.80733 | 0.00052 | 0.04735 |
| ENSMUSG00000038764 | Ptpn3 | 1952.40 | 5.46302 | 1.14505 | 0.00000 | 0.00008 |
| ENSMUSG00000039105 | Atp6v1g1 | 3630.63 | -3.45496 | -0.37411 | 0.00055 | 0.04866 |
| ENSMUSG00000039109 | F13a1 | 422.53 | 4.09417 | 2.00358 | 0.00004 | 0.00934 |
| ENSMUSG00000039304 | Tnfsf10 | 140.30 | 4.21214 | 0.84101 | 0.00003 | 0.00662 |
| ENSMUSG00000039529 | Atp8b1 | 121.41 | 3.70712 | 1.07999 | 0.00021 | 0.02558 |
| ENSMUSG00000039704 | Lmbrd2 | 599.18 | 3.70277 | 0.37714 | 0.00021 | 0.02560 |
| ENSMUSG00000039747 | Orai2 | 19.13 | 3.81439 | 2.01716 | 0.00014 | 0.01987 |
| ENSMUSG00000039987 | Phtf2 | 4103.37 | 3.69012 | 0.27355 | 0.00022 | 0.02647 |
| ENSMUSG00000040276 | Pacsin1 | 10.56 | 3.72924 | 2.98745 | 0.00019 | 0.02424 |
| ENSMUSG00000040283 | Btnl9 | 66.85 | 4.67955 | 1.56367 | 0.00000 | 0.00152 |
| ENSMUSG00000040552 | C3ar1 | 46.70 | 4.05439 | 1.83881 | 0.00005 | 0.01045 |
| ENSMUSG00000040724 | Kcna2 | 125.29 | -3.81865 | -0.89002 | 0.00013 | 0.01987 |
| ENSMUSG00000040820 | Hlcs | 227.28 | 4.40546 | 0.57784 | 0.00001 | 0.00397 |
| ENSMUSG00000041193 | Pla2g5 | 72.97 | -3.79037 | -1.71234 | 0.00015 | 0.02072 |
| ENSMUSG00000041479 | Syt15 | 66.89 | 4.08511 | 1.29319 | 0.00004 | 0.00957 |
| ENSMUSG00000042595 | Fam199x | 248.42 | 3.51044 | 0.58763 | 0.00045 | 0.04234 |
| ENSMUSG00000042828 | Trim72 | 4001.78 | 8.94098 | 0.83880 | 0.00000 | 0.00000 |
| ENSMUSG00000043535 | Setx | 1036.06 | 5.44773 | 0.57750 | 0.00000 | 0.00008 |
| ENSMUSG00000044786 | Zfp36 | 212.61 | 3.98064 | 1.28777 | 0.00007 | 0.01329 |
| ENSMUSG00000045545 | Krt14 | 54.45 | 3.46979 | 7.11309 | 0.00052 | 0.04735 |
| ENSMUSG00000046447 | Camk2n1 | 134.55 | -4.10998 | -0.87477 | 0.00004 | 0.00913 |
| ENSMUSG00000047592 | Nxpe5 | 46.90 | 3.57745 | 2.05964 | 0.00035 | 0.03478 |
| ENSMUSG00000047793 | Sned1 | 524.12 | 3.86728 | 0.90788 | 0.00011 | 0.01786 |
| ENSMUSG00000048490 | Nrip1 | 909.26 | 4.12695 | 1.04233 | 0.00004 | 0.00875 |
| ENSMUSG00000049303 | Syt12 | 52.18 | 3.65316 | 1.36483 | 0.00026 | 0.02875 |

| | | | | | | |
|---|---|---|---|---|---|---|
| ENSMUSG00000051149 | Adnp | 121.64 | 3.81581 | 1.01792 | 0.00014 | 0.01987 |
| ENSMUSG00000051367 | Six1 | 1318.23 | -3.55428 | -0.36399 | 0.00038 | 0.03747 |
| ENSMUSG00000052299 | Ltn1 | 725.92 | 3.62001 | 0.33379 | 0.00029 | 0.03152 |
| ENSMUSG00000052430 | Bmpr1b | 153.05 | 3.54323 | 0.72818 | 0.00040 | 0.03865 |
| ENSMUSG00000052821 | Cysltr1 | 46.20 | 3.91807 | 1.37947 | 0.00009 | 0.01553 |
| ENSMUSG00000052837 | Junb | 220.46 | 4.69446 | 1.55115 | 0.00000 | 0.00146 |
| ENSMUSG00000052920 | Prkg1 | 851.30 | 5.00412 | 0.88929 | 0.00000 | 0.00041 |
| ENSMUSG00000052974 | Cyp2f2 | 45.44 | -4.26231 | -2.23393 | 0.00002 | 0.00588 |
| ENSMUSG00000053113 | Socs3 | 251.37 | 5.04352 | 2.53012 | 0.00000 | 0.00037 |
| ENSMUSG00000053716 | Dusp7 | 282.42 | 3.88792 | 0.91166 | 0.00010 | 0.01696 |
| ENSMUSG00000055065 | Ddx17 | 2670.38 | -4.92401 | -0.37534 | 0.00000 | 0.00054 |
| ENSMUSG00000055489 | Ano5 | 3739.35 | 4.96095 | 0.55222 | 0.00000 | 0.00047 |
| ENSMUSG00000056608 | Chd9 | 657.38 | 4.52158 | 0.48003 | 0.00001 | 0.00275 |
| ENSMUSG00000062593 | Lilrb4a | 7.68 | 3.68259 | 5.63279 | 0.00023 | 0.02705 |
| ENSMUSG00000064357 | ATP6 | 18914.57 | -3.78624 | -0.56950 | 0.00015 | 0.02072 |
| ENSMUSG00000066026 | Dhrs3 | 264.94 | -3.63588 | -0.72395 | 0.00028 | 0.03007 |
| ENSMUSG00000068114 | Ccdc134 | 103.70 | -3.92282 | -0.84049 | 0.00009 | 0.01553 |
| ENSMUSG00000071713 | Csf2rb | 168.55 | 5.85781 | 1.85809 | 0.00000 | 0.00001 |
| ENSMUSG00000071714 | Csf2rb2 | 124.50 | 5.01877 | 1.70365 | 0.00000 | 0.00040 |
| ENSMUSG00000075602 | Ly6a | 711.60 | 3.96812 | 0.78992 | 0.00007 | 0.01353 |
| ENSMUSG00000078816 | Prkcg | 24.96 | -3.96527 | -1.41017 | 0.00007 | 0.01353 |
| ENSMUSG00000079037 | Prnp | 1557.59 | -4.52738 | -0.44792 | 0.00001 | 0.00275 |
| ENSMUSG00000079173 | Zan | 18.14 | 3.91773 | 3.60268 | 0.00009 | 0.01553 |
| ENSMUSG00000081752 | ENSMUSG00000081752 | 20.82 | 5.06835 | 19.81587 | 0.00000 | 0.00035 |
| ENSMUSG00000094013 | ENSMUSG00000094013 | 8.16 | -3.80588 | -5.82147 | 0.00014 | 0.01987 |
| ENSMUSG00000094370 | ENSMUSG00000094370 | 5.40 | -3.70459 | -6.07852 | 0.00021 | 0.02560 |
| ENSMUSG00000095298 | ENSMUSG00000095298 | 12.14 | -3.61415 | -2.36720 | 0.00030 | 0.03191 |
| ENSMUSG00000095752 | ENSMUSG00000095752 | 8.16 | -3.80588 | -5.82147 | 0.00014 | 0.01987 |
| ENSMUSG00000100658 | F730311O21Rik | 9.83 | -3.44325 | -2.18991 | 0.00057 | 0.04934 |
| ENSMUSG00000101249 | ENSMUSG00000101249 | 16141.86 | -6.43893 | -1.07257 | 0.00000 | 0.00000 |
| ENSMUSG00000103560 | ENSMUSG00000103560 | 15.00 | 3.67136 | 2.41636 | 0.00024 | 0.02761 |
| ENSMUSG00000114019 | ENSMUSG00000114019 | 249.45 | -3.44012 | -0.44979 | 0.00058 | 0.04962 |
| ENSMUSG00000114968 | A630019I02Rik | 68.84 | 3.44689 | 0.83125 | 0.00057 | 0.04934 |
| ENSMUSG00000116275 | Zc3h11a | 1100.43 | -4.31656 | -0.68208 | 0.00002 | 0.00498 |
